# Differential roles of delta and theta oscillations in understanding semantic gist during natural audiovisual speech perception: Functional and anatomical evidence

**DOI:** 10.1101/2022.06.21.497061

**Authors:** Hyojin Park, Robin A. A. Ince, Joachim Gross

## Abstract

Understanding the main topic of naturalistic speech in a multi-speaker environment is demanding though the availability of visual speech can be beneficial for speech comprehension. Recent studies provided evidence that low-frequency brain rhythms play an important role in the processing of acoustic speech features. However, at present, the neural dynamics of brain rhythms implementing a higher-order semantic system during naturalistic audiovisual speech perception is unknown. Here we investigated information processing carried by low-frequency oscillations in delta and theta bands for audiovisual speech integration for high-level semantic gist processing using a representational interaction approach. By manipulating the degree of high-level semantic content (speech chunks with high versus low topic probability) using Latent Dirichlet Allocation (LDA) topic modelling algorithm and complexity of speaker environment (single versus multi-speaker), we first found that delta and theta phase exert distinctive roles in high-level semantic processing where delta phase represents auditory and visual inputs synergistically whereas theta band does so redundantly. Next, we show both forms of representational interaction are observed to be greater for speech with low semantic gist, supported by speech comprehension and white matter tractography. Furthermore, we show that the delta phase-specific synergistic interaction in the right auditory, temporal, and inferior frontal areas is sensitive to the speaker environment, whereas theta band activity showing redundant representations is sensitive to semantic content. Our results shed new light on dynamic neural mechanisms of implementing higher-order semantic systems through representational interactions between audiovisual speech information and differential roles of delta and theta bands depending on the speaker environment.

## Introduction

Behavioural and neural evidence on dynamic multi-modal sensory information processing during naturalistic audiovisual speech perception in different speaker environments (single or multi-speaker) has been accumulated in recent years (Park et al., 2016; Hauswald et al., 2018; Park et al., 2018b; Biau et al., 2021; Har-Shai Yahav and Zion Golumbic, 2021). The information processing of different sensory modalities and their integration mechanism has been studied at different levels of speech features, such as acoustic and linguistic levels such as phonemes (Daube et al., 2019; O’Sullivan et al., 2021), lexico-semantics (Broderick et al., 2018) and even during silent lip-reading (Hauswald et al., 2018; Biau et al., 2021; Nidiffer et al., 2021) and occlusion of lip movements (Haider et al., 2022). Low-frequency neural oscillations, which correspond to the temporal structures of units of speech, are known to be prominent in this processing. For example, we have shown that theta phase entrainment in the visual cortex to lip movement (vertical aperture) modulates auditory detection performance during silent speech perception (Biau et al., 2021). However, the role of low-frequency brain rhythms underlying how our brain extracts high-level semantic gist during naturalistic audiovisual speech perception remains largely unknown. In our previous study, we have demonstrated that the neural mechanisms underlying higher-order semantic systems above word-level, such as topic processing during natural speech perception, can be mapped (Park and Gross, 2022).

In the current study, we have scrutinized the functional role of low-frequency oscillations in the processing and representation of multisensory information with regard to the processing of salient semantic gist. Speech chunks were grouped according to their topic probability - high versus low topic probability conditions - from a Latent Dirichlet Allocation (LDA) modelling analysis in order to investigate the difference in neural information regulated by the level of semantic gist across the speech. Corresponding brain signals were segmented along with the speech chunks into the high and low topic probability conditions. Then, the interaction between audio and visual speech information represented in the brain was examined using the representational interactions technique, Interaction Information (McGill, 1954) (also known as Co-Information) based on Information Theory (Shannon, 1948). Representational interaction techniques such as Interaction Information and Partial Information Decomposition (PID) (Ince, 2017; Park et al., 2018b) can determine information representation carried by entrained oscillations either redundantly or synergistically responding to audio and visual speech information. Interaction Information quantifies the difference between the Mutual Information (MI) when the two modalities are observed together and the sum of the MI when each modality is considered alone, which determines redundancy and synergy with opposite signs (see Methods). Redundant, or shared, representation indicates what is learned about the neural response from the visual speech is already obtained from observation of the auditory speech, while synergistic representation indicates that the two modality speech stimuli provide a better prediction when they are considered together than would be expected from observing each individually (for more details, please see Park et al. (2018b)) and recently it has been shown as a robust tool to study neurocognitive architecture at more general level navigating the trade-off between robustness and integration in the brain connectivity (Luppi et al., 2022).

Delta and theta band oscillations in speech perception have been reported as principled computations for temporally relevant speech features such as prosody or intonation and syllabic processing, respectively (Giraud and Poeppel, 2012; Gross et al., 2013; Park et al., 2015). However, their distinctive roles have remained mysterious (but see Kayser et al. (2015) and Giroud et al. (2020)), not to mention audiovisual speech integration in association with high-level linguistic features. We hypothesized differential roles of delta and theta bands in processing semantic gist during audiovisual speech integration involving synergistic and redundant (shared) interaction information processing. We first examined representational interactions (redundancy and synergy) in delta (1-3 Hz) and theta (4-7 Hz) bands between high and low topic probability conditions in a multi-speaker environment using a dichotic listening paradigm. Intriguingly, synergistic interaction between auditory and visual speech is carried by delta phase information (but not theta band), whereas redundant interaction is carried by theta phase information (but not delta band). Furthermore, both types of Interaction Information are greater for semantically less salient speech chunks (low topic probability condition), supported by behavioural performance and white matter tractography. To further delineate the functional role of delta and theta bands, we investigated interactions between two factors of speaker environment and topic probability using the two-way analysis of variance (ANOVA). Here we find delta phase information exercising synergistic information processing is sensitive to the speaker environment in the right auditory, temporal, and inferior frontal areas. Whilst, theta band exercising redundant information processing is mainly dependent on semantic content regardless of speaker environment. In effect, our results demonstrate neural oscillatory dynamics of representational interactions of audiovisual speech in the processing of higher-order semantics and disentangle the functional role of delta and theta bands depending on the speaker environment and high-level semantic gist.

## Results

To understand the neural mechanisms underlying topic keyword processing in a natural audiovisual speech with a cocktail party problem, we used Interaction Information (II) which is an Information Theory-based method to quantify representational interactions. We investigated the representational interactions between dynamic audio and visual speech envelope of the delta and theta frequency bands and the same band activities in the brain (Fig. 1).

**Figure 1.**
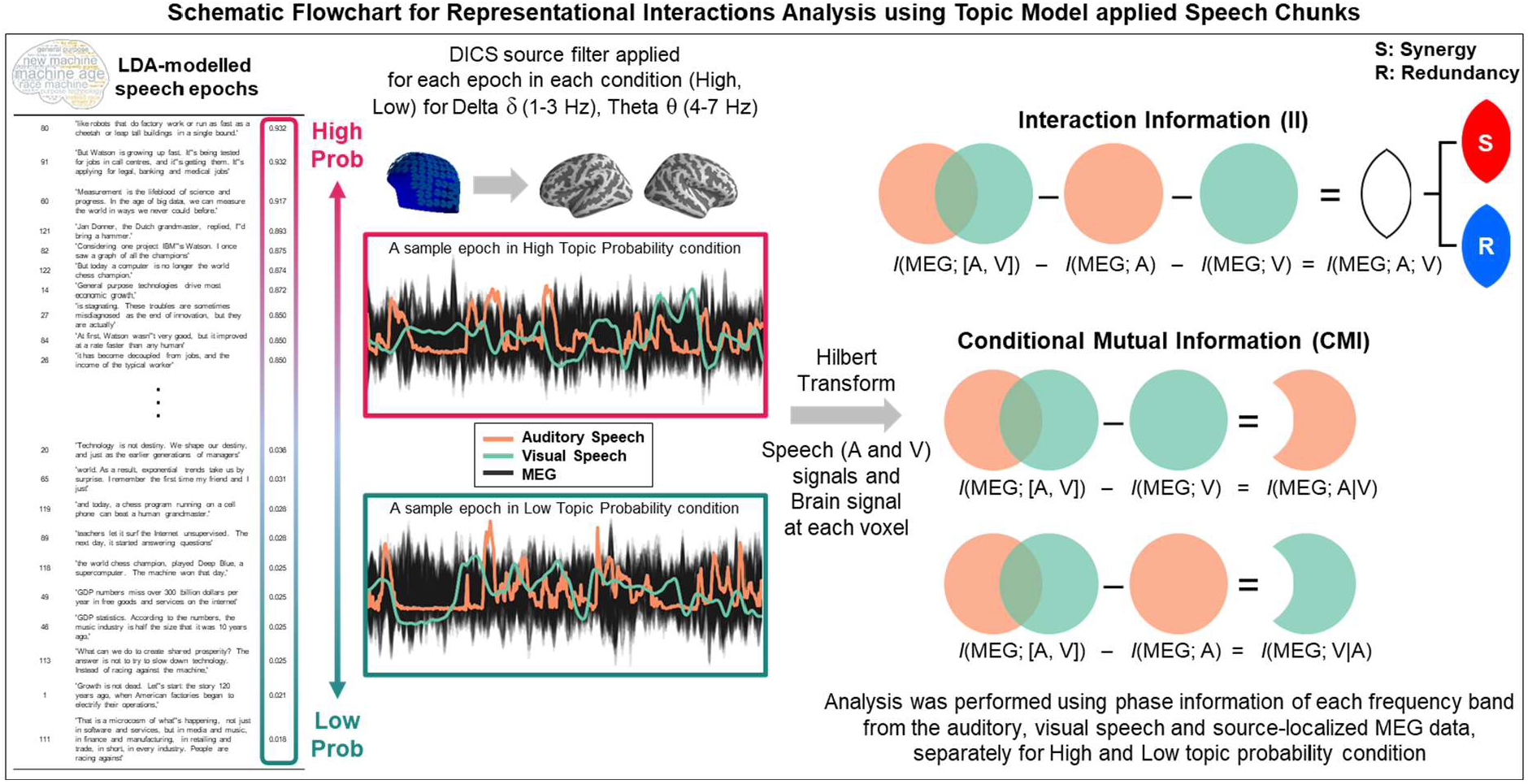
Representational interactions analysis between auditory and visual speech for high and low topic probability conditions. In AV congruent condition, participants heard different auditory speech in each ear with visual speech (face of the speaker) and were instructed to pay attention to one of the auditory talks while ignoring the other. Visual speech (lip movement) was matched to the auditory speech to which the participants paid attention. LDA topic model applied speech chunks were split in to high and low topic probability conditions according to the highest topic cluster (see Park and Gross (2022), which represents the main idea of the talk. MEG data, auditory speech envelope and visual speech signal were band-pass filtered to delta (1-3 Hz) and theta (4-7 Hz) bands, and the DICS source filters were applied to the MEG data. For each speech chunk, Interaction Information (II) and Conditional Mutual Information (CMI) were computed between the phase of auditory speech envelope, visual speech signal, and brain signal at each voxel and averaged across the speech chunks within each condition. CMI computes mutual information between two variables, conditioning out the effect of the third variable. CMI for auditory speech (right middle row), I(MEG; A|V) and CMI for visual speech (right bottom row), I(MEG; V|A) were computed. Interaction Information quantifies the amount of information shared among a set of variables. From the methods for our computation, the outcome provides synergy as positive and redundancy as negative values.

### Modality-specific information for high versus low topic probability characterized by distinctive roles in delta and theta phase information

First, we asked a question if unique modality representations (e.g., representations of auditory speech that are not available from visual speech and vice versa) are modulated by topic probability. To address this question, we investigated the difference between topic probability conditions (high versus low) in a modality-specific manner using CMI (Conditional Mutual Information). For each condition, CMI for auditory speech (right middle row in Fig. 1) was computed between MEG response and auditory speech conditioning out the effect of visual speech, I(MEG; A|V) and CMI for visual speech (right bottom row in Fig. 1) between MEG response and visual speech conditioning out the effect of auditory speech, I(MEG; V|A). Then, each CMI was compared between high and low topic probability conditions.

For CMI for auditory speech (Fig. 2a), I(MEG; A|V), we found a significant difference between high versus low topic probability conditions in the delta band phase, but not in the theta band, in left motor/premotor (BA 4/6), extensive frontal areas including inferior/middle/superior frontal, dorsolateral prefrontal areas (BA 44/46), auditory association cortex, superior/middle/inferior temporal cortices (BA 22), superior/inferior parietal lobule (BA 7) and right inferior frontal, dorsolateral prefrontal areas. For CMI for visual speech (Fig. 2b), I(MEG; V|A), we found a significant difference in the theta band but not in the delta band. Interestingly, CMI was stronger for low topic probability condition in left subcentral, premotor areas (BA 43, 6) and superior/middle/inferior temporal cortices (BA 22). However, the CMI was stronger for high topic probability condition in the left superior/inferior parietal lobule (BA 7), right middle/inferior frontal areas, insular and frontal operculum, primary auditory cortex (BA 41/42), and temporal pole (BA 38). This means that in these areas the representation of modality-specific information is modulated by topic probability.

**Figure 2.**
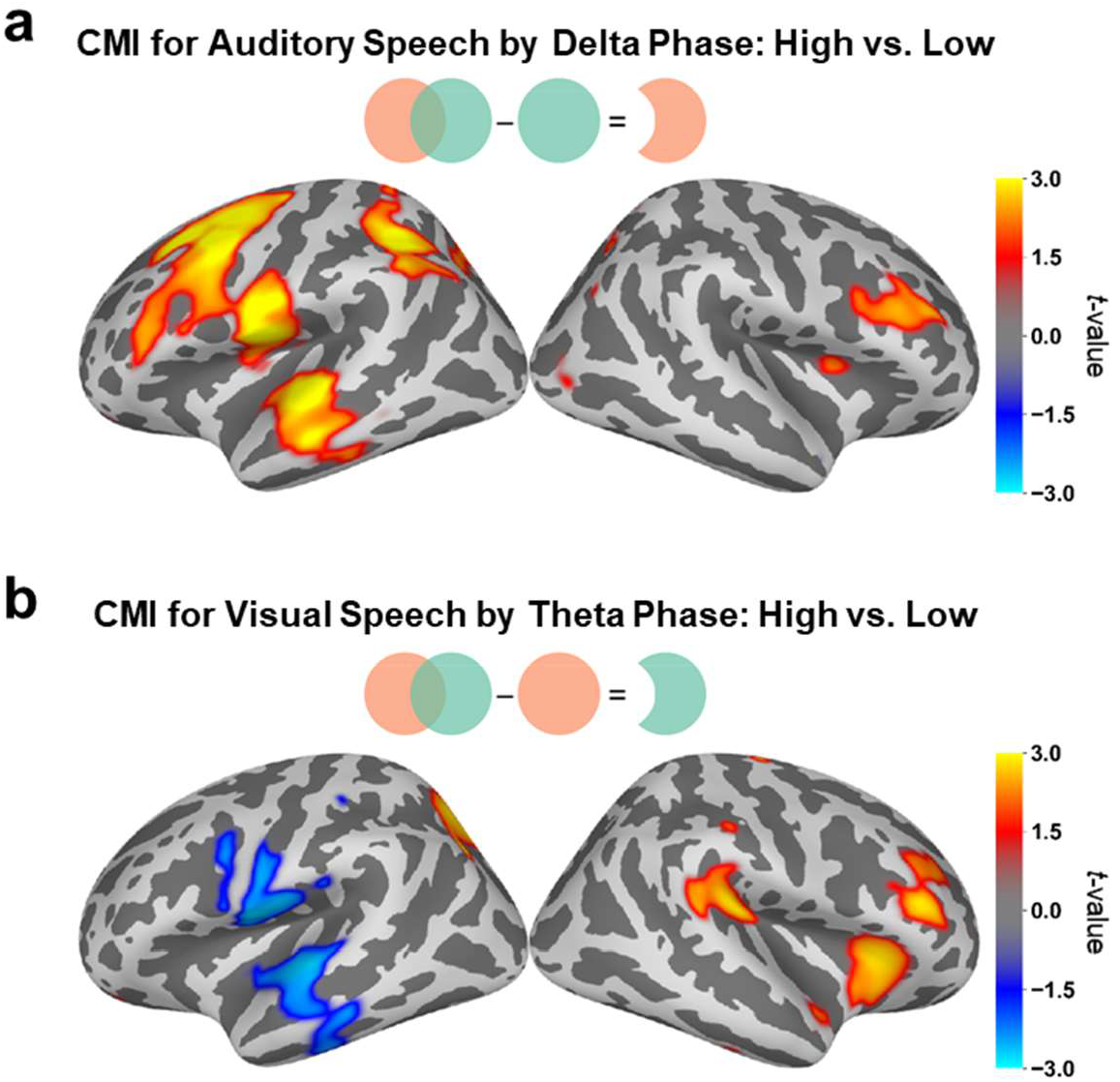
Conditional Mutual Information (CMI) for each modality speech. **a**, CMI difference between high and low probability condition for auditory speech carried by delta phase (but not theta phase; we performed the same analysis in the theta band but found no significant clusters for the semantic contrast). Auditory speech specific information was better represented for the semantically salient speech chunks in left motor/premotor (BA 4/6), extensive frontal areas including inferior/middle/superior frontal, dorsolateral prefrontal areas (BA 44/46), auditory association cortex, superior/middle/inferior temporal cortices (BA 22), superior/inferior parietal lobule (BA 7) and right inferior frontal, dorsolateral prefrontal areas. **b**, CMI difference for visual speech carried by theta phase (but not delta phase; we performed the same analysis in the delta band but found no significant clusters for the semantic contrast). Visual speech specific information is represented in the left superior/inferior parietal lobule (BA 7), right middle/inferior frontal areas, insular and frontal operculum, primary auditory cortex (BA 41/42), temporal pole (BA 38) for semantically salient speech chunks whereas in left subcentral, premotor areas (BA 43, 6) and superior/middle/inferior temporal cortices (BA 22) for semantically less salient speech chunks. Cluster-level permutation test was performed (p < 0.05; two-tailed; 1024 permutations).

### Greater Interaction Information for speech chunks with low semantic gist supported by speech comprehension performance

Next, we assessed the difference between high and low topic probability conditions for each Interaction Information outcome – redundant and synergistic information processing. Intriguingly, the significant topic difference with regards to the synergistic interaction was found in the delta band phase (Fig. 3), not in the theta band, whereas the topic difference with regards to the redundant interaction between auditory and visual speech information was found in theta band phase (Fig. 4), not in the delta band. From the Interaction Information (II) computation, *I*(MEG; [A, V]) - *I*(MEG; A) - *I*(MEG; V), positive Interaction Information outcome values represent synergy and negative outcome values represent redundancy. We first show a grand-average map of Interaction Information (II) for each condition (high and low topic probability) to identify positive and negative outcome values of the effect (panel a in Fig. 3 and Fig. 4). Then compared the conditions statistically using cluster-level permutation (p < 0.05; panel b in Fig. 3 and Fig. 4). To further assess the behavioural relevance of the effect, we performed the regression analysis using speech comprehension accuracy (FDR-corrected, p < 0.05; panel c in Fig. 3 and Fig. 4) to explore the brain regions predicted by the behavioural performance.

**Figure 3.**
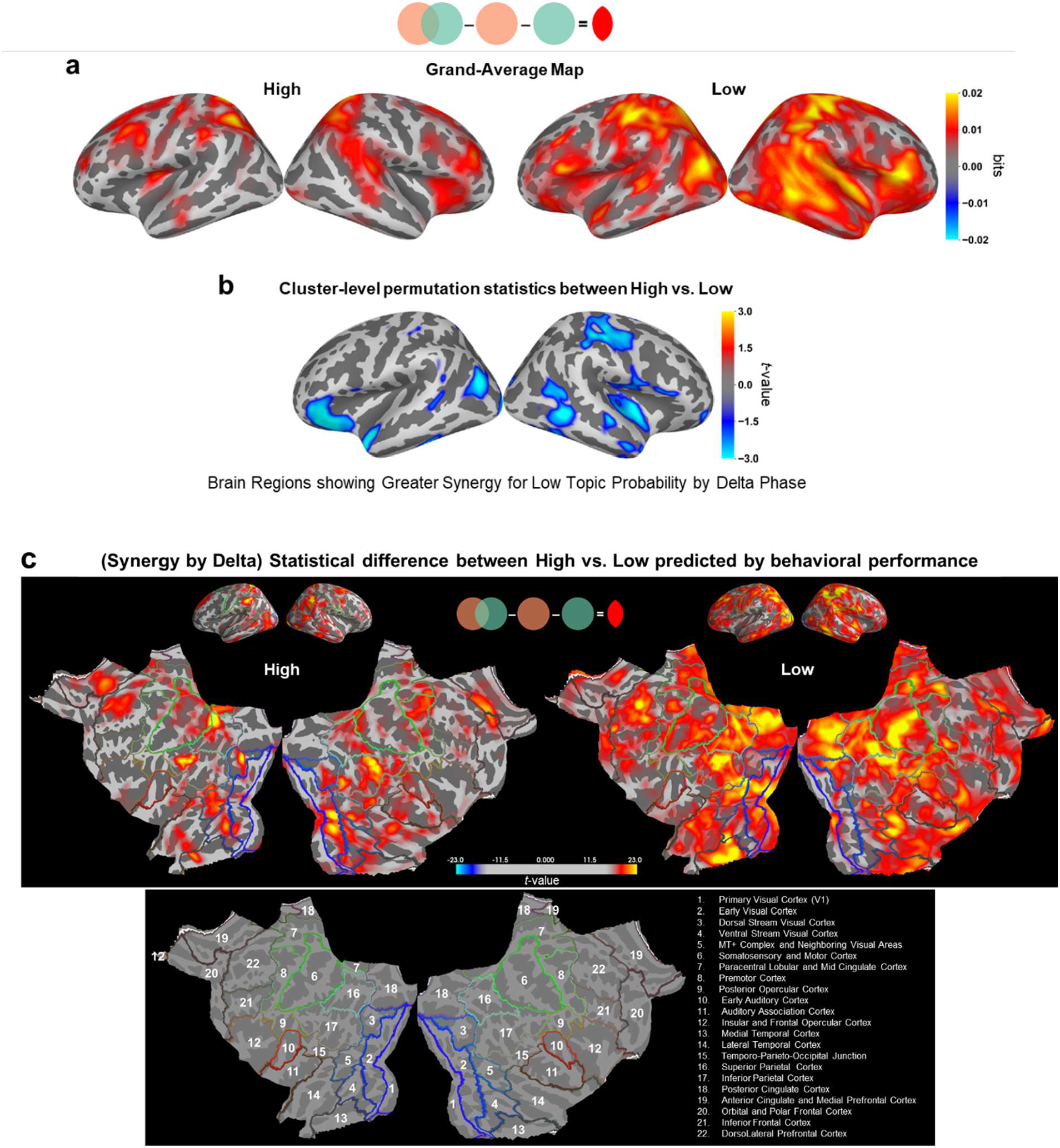
Greater synergistic interactions between auditory and visual speech for speech with low semantic gist carried by delta phase information. **a**, Grand-average map of values from Interaction Information computation for each high and low topic probability condition. Both conditions show positive values indicating synergistic interactions. **b**, Statistical comparison between high versus low topic probability conditions was performed using cluster-level permutation (p < 0.05; 1024 permutations). Brain regions shown in blue indicate greater synergy for low topic probability condition carried by delta phase information. **c**, Regression analysis using speech comprehension accuracy (FDR-corrected, p < 0.05). In the linear regression model, speech comprehension scores from all subjects were used as a regressor. Results show a strong speech comprehension prediction effect for low topic probability condition by delta phase synergy in the left middle temporal, dorsolateral prefrontal, superior/inferior parietal, early visual cortices and the right motor, somatosensory, lateral temporal, superior parietal, posterior cingulate cortices. Results are shown in classic (top) as well as flat brain templates. For the results using the flat template, please see the region information from the HCP-MMP1.0 combined atlas given below.

**Figure 4.**
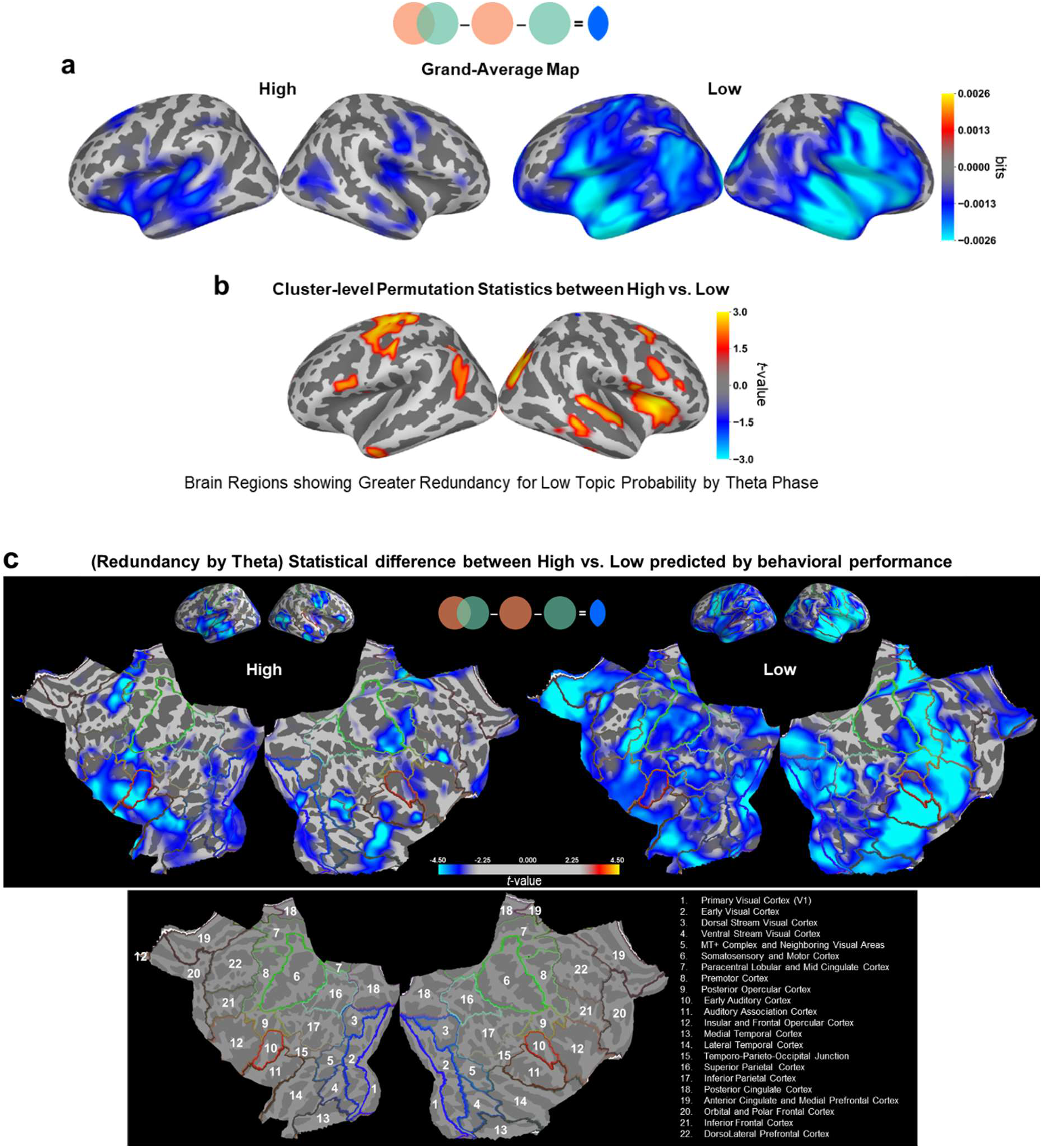
Greater redundant interactions between auditory and visual speech for speech with low semantic gist carried by theta phase information. **a**, Grand-average map of values from Interaction Information computation for each high and low topic probability condition. Both conditions show negative values indicating redundant interactions. **b**, Statistical comparison between high versus low topic probability conditions was performed using cluster-level permutation (p < 0.05; 1024 permutations). Brain regions shown in red indicates greater redundancy for low topic probability condition carried by theta phase information. **c**, Regression analysis using speech comprehension accuracy (FDR-corrected, p < 0.05). In the linear regression model, speech comprehension scores from all subjects were used as a regressor. Results show a strong speech comprehension prediction effect for low topic probability condition by theta phase redundancy in bilateral frontal, temporal and somatosensory areas with greater right hemisphere engagement. Results are shown in classic (top) as well as flat brain templates. For the results using the flat template, please see the region information from the HCP-MMP1.0 combined atlas given below.

Interestingly, Interaction Information was shown to be greater for low topic probability condition for both redundancy by theta phase and synergy by delta phase. The synergistic interactions (positive values) between audio and visual speech features found in the delta band phase was stronger for low topic probability condition as well in extensive bilateral parieto-occipital and right temporal areas. The significant difference between the conditions (Fig. 3b) was mapped on to the left inferior frontal, superior temporal pole (BA 38), parieto-occipital areas (BA 39/19), and right motor (BA 4), somatosensory (BA 1/2), inferior frontal (BA 44/45), superior/middle temporal sulcus (BA 22/37).

Note that the statistical map (high versus low contrast) was cool colour-coded due to the positive values indicating synergistic representation resulting in greater response for low topic probability condition. When each synergy map for high and low topic probability conditions was predicted by speech comprehension accuracy (Fig. 3c), the patterns were similar to the grand-average map (Fig. 3a) in terms of stronger patterns for low topic probability condition. Results show strong behavior (speech comprehension) prediction effect for low topic probability condition by delta phase synergy in the left middle temporal, dorsolateral prefrontal, superior/inferior parietal, early visual cortices and the right motor, somatosensory, lateral temporal, superior parietal, posterior cingulate cortices.

The redundant representation (negative values) between audio and visual speech features found in the theta band was stronger for low topic probability condition in extensive bilateral frontal, temporal, and somatosensory areas. The significant difference between the conditions (Fig. 4b) was localized in the left motor and premotor cortices (BA 4/6), inferior frontal area (BA 44), posterior superior temporal sulcus (BA 39), and right insular and frontal opercular cortex, auditory association cortex, visual areas (BA 18/19). Note that the statistical map (high versus low contrast) was warm colour-coded due to the negative values indicating redundant representation resulting in greater response for low topic probability condition. When each redundancy map for high and low topic probability conditions was predicted by speech comprehension accuracy (Fig. 4c), the patterns were similar to the grand-average map (Fig. 4a). Results show strong behavior (speech comprehension) prediction effect for low topic probability condition by theta phase redundancy in bilateral frontal, temporal and somatosensory areas with greater right hemisphere engagement. For high topic probability condition, the left temporal areas, including the anterior portion and dorsolateral prefrontal cortex, and the right inferior frontal and middle/inferior temporal cortices were shown to predict the speech comprehension score.

### Anatomical evidence supports differential patterns of Interaction Information by delta and theta for speech with low semantic gist

Intrigued by the dissociated pattern of interaction information - redundant representation in theta band whereas synergistic representation in delta band - in low topic probability condition, we hypothesized that stronger interaction information for speech chunks with low topic probability condition, characterized by low semantic core, is supported by stronger anatomical connectivity. We performed probabilistic tractography analysis using diffusion-weighted images from the same subjects in the FSL (using modules BEDPOSTX and PROBTRACKX). Five major tracts - Inferior Fronto-Occipital Fasciculus, Inferior Longitudinal Fasciculus, Superior Longitudinal Fasciculus, Arcuate Fasciculus (also known as Superior Longitudinal Fasciculus Temporal Part) and Uncinate Fasciculus in the left hemisphere with 2 mm resolution - were used as seed regions for seed-based probabilistic tractography analysis. Outcome values represent connectivity distribution based on Bayesian estimation from the seed voxel (Behrens et al., 2003; Behrens et al., 2007), i.e., the number of samples that pass through the seed voxel. We extracted the volumes (the number of voxels) showing the connectivity with the seed (Fig. 5a). The relationship with Interaction Information was assessed by correlating the extracted volumes from each seed fiber tract with the difference between low and high topic probability conditions normalized by their sum for each frequency band (delta and theta) using Spearman rank-order correlation. We here have diffusion-weighted images for 41 out of 44 subjects; thus, the correlation analysis was performed using 41 subjects’ data. Here we performed the correlational analysis via the regions of interest approach using the HCP-MMP1.0 combined atlas (also shown in the inset at the bottom of Fig. 3c and 4c) from HCP-MMP1.0 (Human Connectome Project Multi-Modal Parcellation version 1.0) (Glasser et al., 2016).

**Figure 5.**
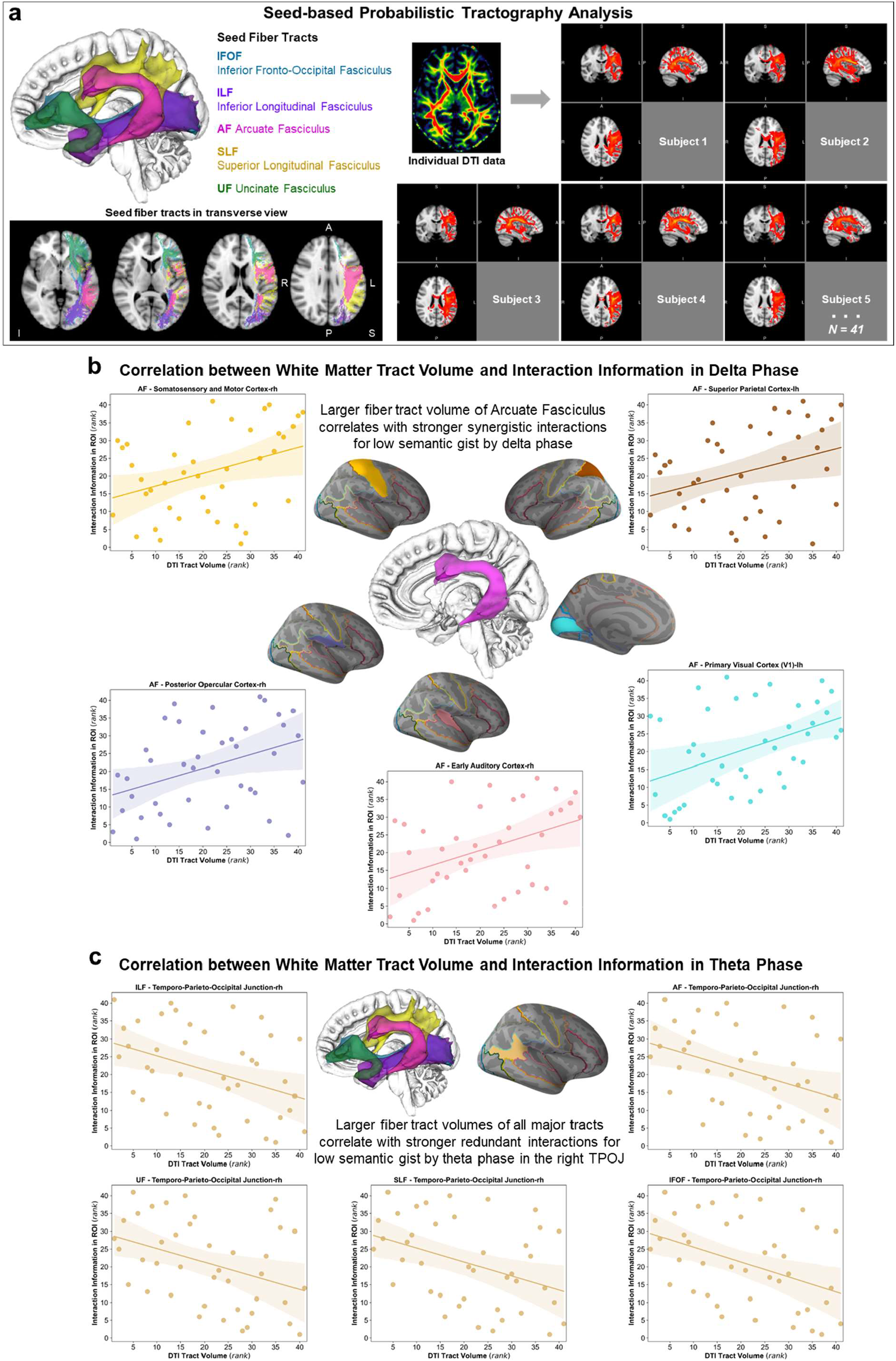
Seed-based probabilistic tractography analysis using five major white matter tracts supports the evidence for stronger Interaction Information for low topic probability condition. **a, Seed-based probabilistic tractography analysis using diffusion-weighted images from the same subjects was performed in FSL.** Five major tracts - Inferior Fronto-Occipital Fasciculus, Inferior Longitudinal Fasciculus, Superior Longitudinal Fasciculus, Arcuate Fasciculus (also known as Superior Longitudinal Fasciculus Temporal Part) and Uncinate Fasciculus in the left hemisphere with 2 mm resolution - were used as seed regions. Volumes (number of voxels) with connectivity distribution from the seed fiber tracts were extracted. The relationship with Interaction Information was assessed by correlating the extracted volumes from each seed fiber tract with the difference between low and high topic probability conditions normalized by their sum for each frequency band (delta and theta) using Spearman rank-order correlation. The Interaction Information values were extracted from each region of interest in the HCP-MMP1.0 combined atlas. The left hemisphere is shown on the right. *Abbreviations. S*: Superior; *I*: Inferior; *A*: Anterior; *P*: Posterior; *L*: Left; *R*: Right. **b, White matter volumes and delta phase information**. Larger fiber tract volume of Arcuate Fasciculus correlate with stronger synergistic interactions by delta phase in the right early auditory cortex (r = 0.40, p = 0.01), posterior opercular cortex (r = 0.35, p = 0.023), somatosensory and motor cortex (r = 0.36, p = 0.023), left primary visual cortex (r = 0.43, p = 0.006), and superior parietal cortex (r = 0.32, p = 0.039). Significant positive correlation represents the relationship with synergistic interactions. **c, White matter volumes and theta phase information**. Larger fiber tract volumes of all the five major tracts (shown in an inset on the middle left) correlate with stronger redundant interactions by theta phase in the right temporo-parieto-occipital junction (TPOJ; shown in an inset on the middle right). Correlation with Inferior Fronto-Occipital Fasciculus (r = -0.42, p = 0.007), Inferior Longitudinal Fasciculus (r = -0.39, p = 0.012), Arcuate Fasciculus (r = -0.38, p = 0.013), Superior Longitudinal Fasciculus (r = -0.39, p = 0.011) and Uncinate Fasciculus (r = -0.38, p = 0.015). Significant negative correlation represents the relationship with redundant interactions.

The results show that larger fiber tract volume of Arcuate Fasciculus correlate with stronger synergistic interactions by delta phase in the following brain regions (Fig. 5b): right early auditory cortex (r = 0.40, p = 0.01), posterior opercular cortex (r = 0.35, p = 0.023), somatosensory and motor cortex (r = 0.36, p = 0.023), left primary visual cortex (r = 0.43, p = 0.006), and superior parietal cortex (r = 0.32, p = 0.039). Since the positive values of the Interaction Information indicate synergistic interactions, the significant positive correlation represents the relationship with synergistic interactions.

For theta phase, however, the results show different patterns that larger fiber tract volumes of all the five major tracts correlate with stronger redundant interactions by theta phase in the right temporo-parieto-occipital junction as follows (Fig. 5c): with Inferior Fronto-Occipital Fasciculus (r = -0.42, p = 0.007), Inferior Longitudinal Fasciculus (r = -0.39, p = 0.012), Arcuate Fasciculus (r = -0.38, p = 0.013), Superior Longitudinal Fasciculus (r = -0.39, p = 0.011) and Uncinate Fasciculus (r = -0.38, p = 0.015). Since the negative values of the Interaction Information indicate redundant interactions, the significant negative correlation represents the relationship with redundant interactions.

### Greater interactions for low semantic gist: Synergy by delta sensitive to the speaker condition, whereas redundancy by theta for tracking contents

Interestingly redundant and synergistic interactions between auditory and visual speech were identified stronger for low topic probability condition. We hypothesized that greater interaction information processing for speech with less salient semantic gist might be due to recompense for improving the level of speech comprehension at behavioural performance in a multi-speaker situation. Thus, we further characterize the dynamic information representations by comparing this multi-speaker condition to a single speaker condition, i.e., a diotic listening condition with visual speech for which the Interaction Information analysis was performed as well. We ran a two-way analysis of variance (ANOVA) test with factors of speaker condition (single versus multi-speaker) and topic probability (high versus low topic probability) for delta and theta frequency bands separately. Here we used the same predefined cortical parcellation, HCP-MMP1.0 combined atlas (Human Connectome Project Multi-Modal Parcellation version 1.0) (Glasser et al., 2016) and reported results of auditory (early and association cortex), temporal, frontal areas in both hemispheres. For more detailed results in other brain areas, including visual, parietal, somatosensory and motor regions, please see Supplementary Figure 1.

We here report three main findings from this analysis (Fig. 6): The ANOVA revealed 1) the Interaction Information values of delta band phase in left auditory, temporal, and inferior frontal areas were observed for the main effect of speaker condition (left two columns in the top row in Fig. 6a-b), whereas 2) the Interaction Information values of delta band in right auditory, temporal, frontal areas were observed for the interaction effect of speaker condition and topic probability (right two columns in top row in Fig. 6a-b), and 3) the Interaction Information values of theta band in right auditory cortices and bilateral temporal, frontal areas were observed for the main effect of topic probability (bottom rows in Fig. 6a-b). Interestingly, similar patterns show in other areas, including visual, parietal, somatosensory, motor, and frontal areas displayed in Supplementary Figure 1.

**Figure 6.**
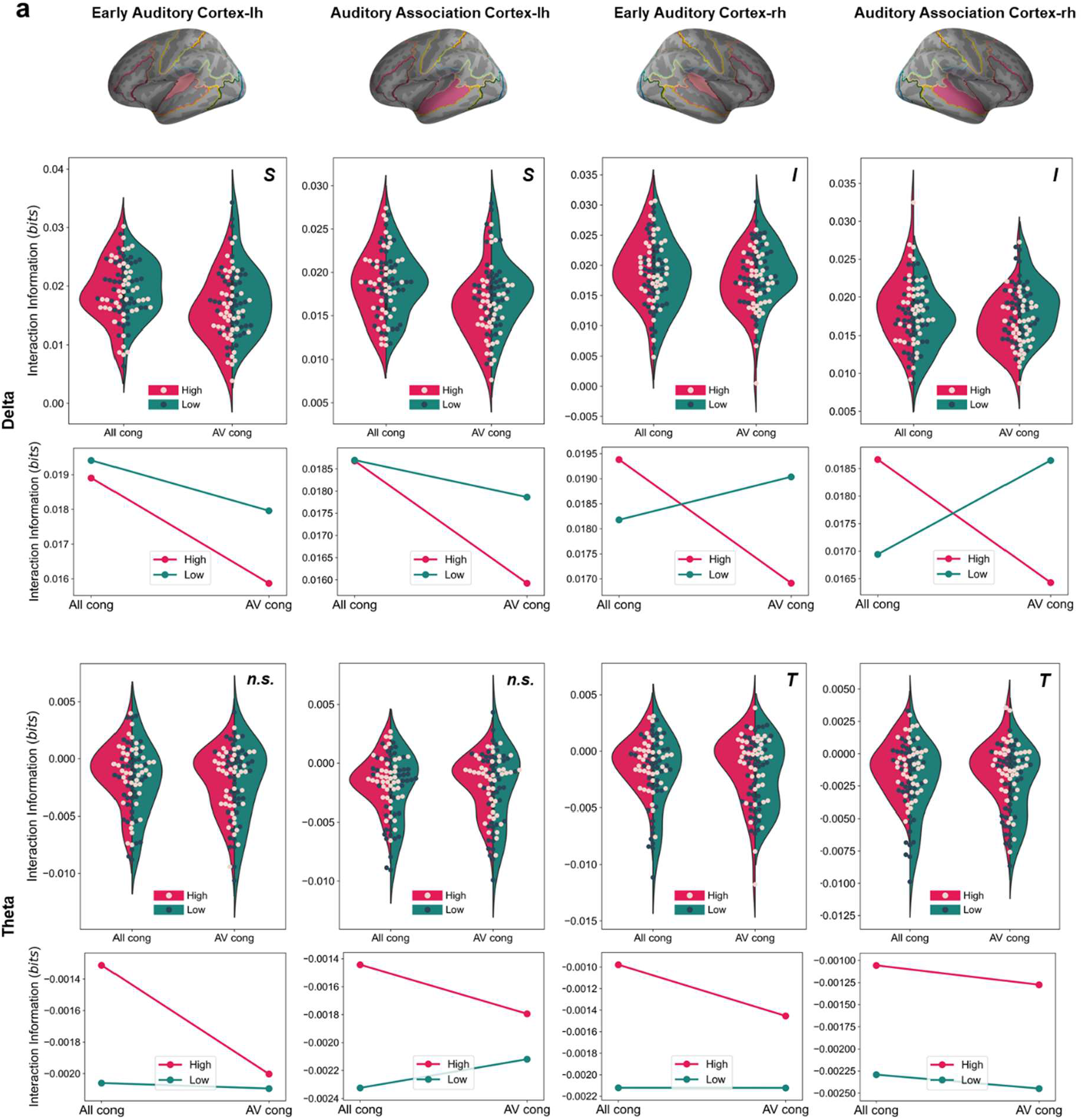
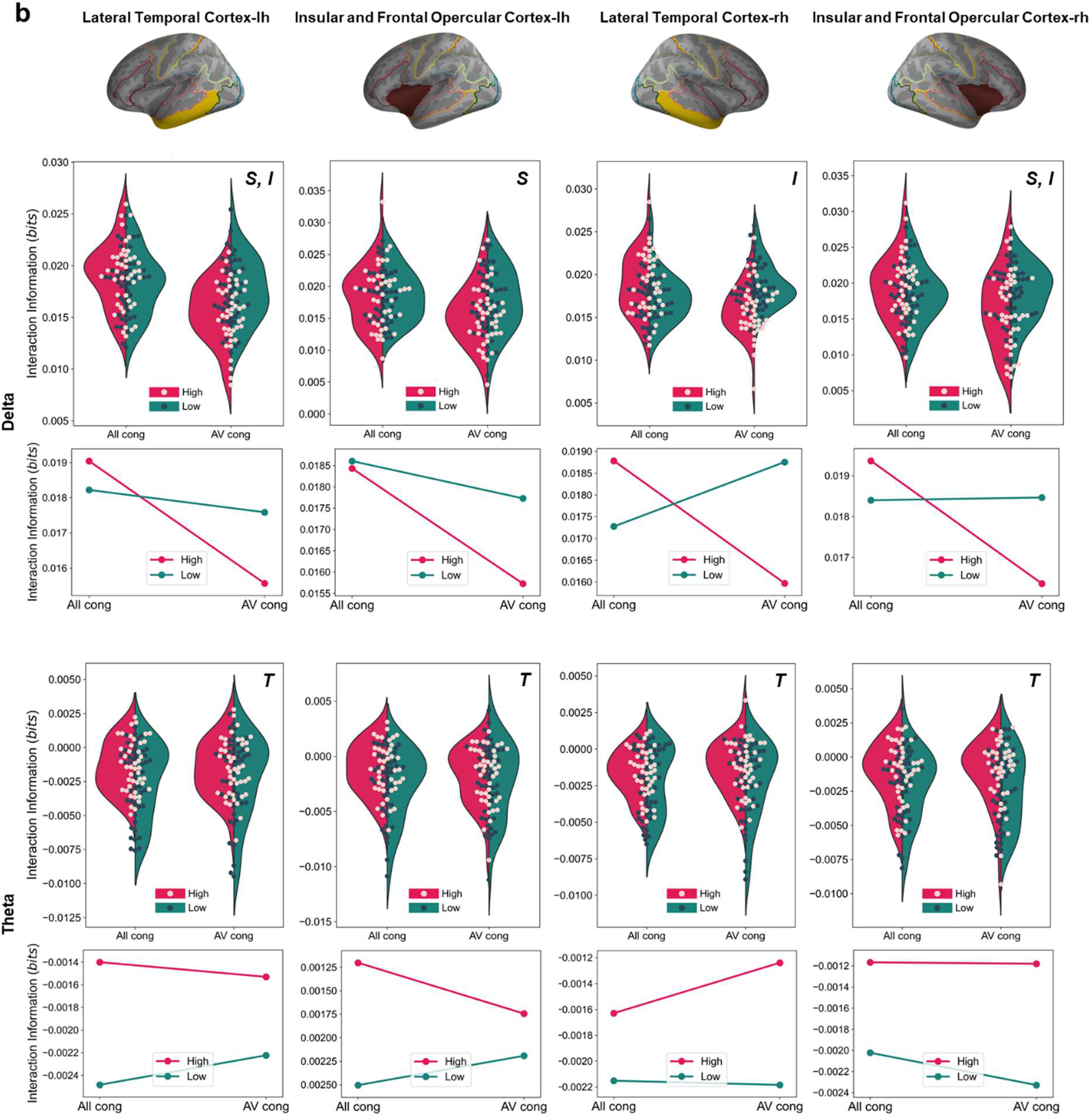
Two-way analysis of variance (ANOVA) test with factors of speaker condition (single versus multi-speaker) and topic probability (high versus low topic probability) for delta and theta band phase Interaction Information. Interaction Information values were extracted from All congruent and AV congruent conditions and high and low topic probability conditions, and two-factor ANOVA was performed. **a**, Results for auditory areas. **b**, Results for lateral temporal and inferior frontal areas. Interaction Information values of delta band phase in left auditory, temporal, and inferior frontal areas show the main effect of speaker condition (marked by *S*), whereas in the same regions in the right hemisphere show interaction effect between speaker condition and topic probability (marked by *I*). Interaction Information values of theta band in the right auditory cortices and bilateral temporal and frontal areas show the main effect of topic probability (marked by *T*). Please see Supplementary Figure 1 for more results from all brain regions, including visual, parietal, somatosensory and motor areas in the atlas.

F- and P-values for each region are as follows: Delta band in the left hemisphere (left two columns in top row in Fig. 6a-b): early auditory cortex (main effect of speaker condition: F_1,172_ = 7.47, p = 0.007), auditory association cortex (main effect of speaker condition: F_1,172_ = 9.26, p = 0.002), lateral temporal cortex (main effect of speaker condition: F_1,172_ = 19.39, p = 2 × 10^−5^; interaction effect of speaker condition and topic probability: F_1,172_ = 9.27, p = 0.003), insular and frontal opercular cortex (main effect of speaker condition: F_1,172_ = 6.45, p = 0.01); Delta band in the right hemisphere (right two columns in top row in Fig. 6a-b): early auditory cortex (interaction effect of speaker condition and topic probability: F_1,172_ = 4.46, p = 0.03), auditory association cortex (interaction effect of speaker condition and topic probability: F_1,172_ = 11.13, p = 0.001), lateral temporal cortex (interaction effect of speaker condition and topic probability: F_1,172_ = 21.70, p = 6 × 10^−6^), insular and frontal opercular cortex (main effect of speaker condition: F_1,172_ = 4.64, p = 0.03; interaction effect of speaker condition and topic probability: F_1,172_ = 5.08, p = 0.02); Theta band in the left hemisphere (left two columns in bottom row in Fig. 6a-b): early auditory cortex (*n.s*.), auditory association cortex (*n.s*.), lateral temporal cortex (main effect of topic probability: F_1,172_ = 5.67, p = 0.01), insular and frontal opercular cortex (main effect of topic probability: F_1,172_ = 4.00, p = 0.04); Theta band in the right hemisphere (right two columns in bottom row in Fig. 6a-b): early auditory cortex (main effect of topic probability: F_1,172_ = 4.16, p = 0.04), auditory association cortex (main effect of topic probability: F_1,172_ = 10.20, p = 0.001), lateral temporal cortex (main effect of topic probability: F_1,172_ = 5.16, p = 0.02), insular and frontal opercular cortex (main effect of topic probability: F_1,172_ = 7.12, p = 0.008). Note that positive Interaction Information values indicate synergistic interactions and negative Interaction Information values indicate redundant interactions, so the main effect for topic probability shown in theta phase results (bottom rows in Fig. 6a-b) depicts greater redundant interactions for low topic probability condition.

## Materials and Methods

### Participants

The study was approved by the local ethics committee at the College of Science and Engineering, University of Glasgow (ethics code: CSE01321) and was conducted in accordance with the ethical guidelines in the Declaration of Helsinki. Forty-six healthy, native English speaking volunteers participated in the study. All participants had normal or corrected-to-normal vision and reported no history of neurologic or psychiatric disorders. They all reported normal hearing (confirmed by two hearing tests using research applications on an iPad: uHear (Unitron Hearing Limited) and Hearing-Check (RNID)) and all right-handed confirmed by Edinburgh Handedness Inventory (Oldfield, 1971). Data from 44 subjects were analyzed (26 females, 18 males; age range: 18-30 years; mean age: 20.54 ± 2.58 years) after two subjects were excluded since one subject fell asleep and one had excessive signal noise. Other analyses of these data were reported in previous publications (Park et al., 2016; Park et al., 2018b). All subjects provided informed written consent before the experiment and received monetary compensation or course credits for their participation.

### Stimuli

#### Naturalistic audiovisual talks

Eleven TED talks (www.ted.com/talks/) were downloaded and modified to be appropriate for our own filming of a professional speaker talking continuously. Transcription for each talk was edited to be appropriate for our own filming by editing words such as referring to visual materials, the gender of the speaker etc. Each talk addresses a specific topic belonging to informative, persuasive, and inspiring categories, albeit they no longer provide these categories explicitly. The contents of the talks were further validated in a separate behavioural study (33 participants with 19 females; aged 18-31 years; mean age: 22.27 ± 2.64 years) in which the participants rated each talk in terms of arousal, familiarity, valence, complexity, significance (informativeness), agreement (persuasiveness), concreteness, self-relatedness, and level of understanding using Likert scale (Likert, 1932) 1 to 5 (for an example of concreteness, 1: very abstract, 2: abstract, 3: neither abstract nor concrete, 4: concrete, 5: very concrete). Eight out of eleven videos were selected for the MEG experiment after three talks with excessive mean scores (below 1 and over 4) were excluded. Selected talks were used in different experimental conditions, which were also used in our previous report (Park et al., 2016).

Furthermore, in order to further rule out the possibility of the neural representations driven by emotional valence, we performed sentiment analysis on speech text materials. Using VADER (Valence Aware Dictionary for sEntiment Reasoning) (Hutto and Gilbert, 2014) model in the NLTK 3.5 (Natural Language Toolkit) package (Bird et al., 2009), which provides both polarities (positive/neutral/negative) and intensity (strength) of emotion. The outcome of the model provides scores for each category of positive, neutral, negative and compound. The maximum values for positive, neutral and negative are 1.0, and the compound score is a normalized value across positive, neutral, and negative scores to be between -1 and 1. The results for talks show near-zero compound scores meaning that the speech materials are all with a neutral sentiment (see Supplementary Table 2 in Park and Gross (2022).

High-quality audiovisual video clips (with a sampling rate of 48 kHz for audio and 25 frames per second (fps) for video in 1,920 × 1,080 pixels) were filmed by a professional filming company while a professional male speaker was speaking each talk that lasted 7 to 9 minutes. Using Final Pro Cut X (Apple, Cupertino, CA), the videos were edited for different experimental conditions (see below) by manipulating combinations of auditory and visual talks in which some conditions involve a dichotic listening paradigm.

#### Experimental tasks

During the MEG measurements, we recorded brain activity from the participants during four different experimental tasks, as reported in (Park et al., 2016). In the current study, we analyzed data from All congruent and AV congruent conditions. In All congruent condition, the auditory talk is presented diotically with the matching video, and the participants were instructed to pay attention to auditory and visual speech (i.e., lip movement) naturally. In AV congruent condition, two different auditory talks were presented to each ear and participants were instructed to pay attention to one of the talks; thus, the other talk serves as a distractor. The visual speech was matched to the auditory speech the participants paid attention to. Here the side of the attention (i.e., left or right) was counterbalanced across participants (22 participants each either to the left or the right side). In this condition, visual speech is beneficial to the understanding of task-relevant auditory speech. In the data analysis, we pooled across both groups so that attentional effects for the left or right side are expected to cancel out. Throughout all the conditions, participants were instructed to pay attention to visual speech and their eye movements were examined using an eye tracker (EyeLink 1000, SR Research). The audiovisual stimuli were presented via Psychtoolbox (Brainard, 1997) in MATLAB R2019b. Auditory stimuli were presented at a 48 kHz sampling rate via a sound pressure transducer through two five-meter-long plastic tubes terminating in plastic insert earpieces, and visual stimuli were delivered at 25 fps (frame per second) as mp4 format with a resolution of 1,280 × 720 pixels.

#### Speech comprehension questionnaire

To assess the level of comprehension of the talk, a questionnaire for speech comprehension was administered after the condition. The questionnaire consists of 10 questions about the talk to which the participants were instructed to pay attention, such as “What is the speaker’s job?” and “What would be the best title of this talk?”. The questionnaire for each talk was validated in a separate behavioral study (see Park et al. (2016); 16 subjects; 13 females; aged 18-23 years; mean age: 19.88 ± 1.71 years) to achieve a similar level of accuracy, response time, and the length of the questionnaire (word count). Participants show high comprehension scores for the two conditions. The mean accuracy (%) was 85 ± 1.66 for All congruent condition and 83.40 ± 1.73 for AV congruent condition. The accuracy did not differ between the two conditions (paired t-test, t_43_ = 0.76, p = 0.45).

### Data acquisition

#### MEG

MEG data were acquired using a 248 magnetometers whole-head MEG system (MAGNES 3600 WH, 4-D Neuroimaging) in a magnetically shielded room at the Centre for Cognitive Neuroimaging (CCNi), University of Glasgow. The sampling rate at the measurement was 1,017 Hz. The signals were then resampled to 250 Hz and denoised with information from the reference sensors using the denoise_pca function in the FieldTrip toolbox (Oostenveld et al., 2011). Bad MEG sensors were excluded using a semi-automatic approach by visual inspection and automatic artifact rejection in Fieldtrip. Electrooculographic (EOG) and electrocardiographic (ECG) artefacts were rejected using independent component analysis (ICA).

#### MRI and DWI

High-resolution anatomical T1-weighted MRI of each participant was acquired at 3 T Siemens Trio Tim scanner (Siemens, Erlangen, Germany) at the Centre for Cognitive Neuroimaging (CCNi), the University of Glasgow, with the following parameters: 1.0 × 1.0 × 1.0 mm^3^ voxels; 192 sagittal slices; field of view (FOV): 256 × 256 matrix. Diffusion-weighted images were acquired with a twice-refocused spin-echo EPI sequence (88 axial slices, TR = 12000 ms, TE = 100 ms, field of view (FOV): 128 × 128 matrix with a voxel size of 1.72 × 1.72 × 1.7 mm^3^). Diffusion-weighted volumes were isotropically distributed along 60 diffusion-encoding gradient directions with a b-value of 1000 s/mm^2,^ and seven non diffusion-weighted volumes (b0) were interspersed for anatomical reference for offline motion correction. DWI images were not acquired from 3 subjects among the 44 subjects; thus the probabilistic tractography analysis results (see below) are obtained from the 41 subjects.

### Data analysis

#### Coregistration between MEG and MRI T1 data

Anatomical T1 MR images recorded from each participant were co-registered to the MEG coordinate system via a semi-automatic approach to map MEG sensor data onto the brain’s source space. Before the MEG recording, fiduciary landmarks - nasion, bilateral preauricular points - were identified, and five head-position indicator coils (HPI coils) were digitized. Over 200 extra points on the scalp were acquired to co-register the MEG source space data with the MRI image. The fiduciary landmarks were also manually identified in the individual’s MR images. Based on these landmarks, both MEG and MRI coordinate systems were initially aligned, followed by numerical optimization achieved by using the ICP algorithm (Besl and McKay, 1992).

#### Source space analysis

MEG results shown on source space were analyzed in Fieldtrip (Oostenveld et al., 2011) and MNE-Python (Gramfort et al., 2014) following the standard procedure for inverse solution in each software. First, a head model was created for each individual from their structural MRI using normalization and segmentation routines in FieldTrip and SPM8. The leadfield was computed using a single-shell volume conductor model (Nolte, 2003) employing an 8 mm grid defined on the MNI (Montreal Neurological Institute) template. For spatial normalization, the template grid was then linearly transformed into individual headspace. In MNE-Python, the forward model construction and MRI segmentation were performed in FreeSurfer (Dale et al., 1999). Boundary Element Model (BEM) for each subject’s MRI was created using the FreeSurfer watershed algorithm, and surface-based source space with a 4.9 mm source spacing resolution was computed. For spatial normalization, the individual source map was morphed to the template MRI, fsaverage.

For the delta and theta frequency band, cross-spectral density matrices were computed using fast Fourier transform on 1 s segments of data after applying multitaper with ±2 Hz frequency smoothing. Then, DICS beamforming algorithm (Gross et al., 2001) was applied for source localization. The beamformer coefficients were computed sequentially for all frequencies from 1 to 7 Hz for the dominant source direction in all voxels with regularization of 7% of the mean across eigenvalues of the cross-spectral density matrix.

#### Probabilistic tractography analysis

Diffusion MRI data were analyzed using the FSL (Smith et al., 2004). As a preprocessing step, the T1-weighted structural scans were skull-stripped and aligned to AC-PC in Talairach space. Using the seven non diffusion-weighted b0 images, eddy current correction (motion and image distortion corrections) parameters for diffusion-weighted images were computed via rigid-body transformations in FDT (FMRIB’s Diffusion Toolbox). The parameters were applied to all 60 diffusion-weighted volumes, and the gradient direction for each volume was corrected using the rotation parameters. Next, seed-based white matter connectivity was analyzed using probabilistic fiber tracking in FDT. Five major white-matter tracts from JHU white-matter tractography atlas (Mori et al., 2005; Hua et al., 2008) were used as seed regions - Inferior Fronto-Occipital Fasciculus, Inferior Longitudinal Fasciculus, Superior Longitudinal Fasciculus, Arcuate Fasciculus (also known as Superior Longitudinal Fasciculus Temporal Part) and Uncinate Fasciculus in the left hemisphere with 2 mm resolution. Then, the fiber orientation was estimated in each voxel by the module BEDPOSTX using a crossing fiber model with up to two directions per voxel. Afterwards, the seed-based probabilistic tractography was computed using the module PROBTRACKX with standard parameters of a curvature threshold of 0.2, a maximum number of steps of 2000, a step length of 0.5 and 5000 streamlines per seed region voxel. PROBTRACKX, probabilistic tracking with crossing fibres, provides connectivity distribution based on Bayesian estimation (Behrens et al., 2003; Behrens et al., 2007). It repeatedly samples from the distributions of voxel-wise principal diffusion directions. Taking repeated streamlined samples, it builds the histogram of the posterior distribution. The output values represent the connectivity distribution from the seed voxel, i.e., the number of samples that pass through the seed voxel. To assess the relationship with Interaction Information measures, we extracted the volumes (the number of voxels) that show the connectivity with the seed. Next, the each of fiber tract volume was correlated with Interaction Information values (low > high topic probability condition normalized by their sum; (II_low_ – II_high_) / (II_low_ + II_high_)) in the each region of interest for each frequency band using Spearman rank-order correlation at p < 0.05 (function: scipy.stats.spearmanr).

Here we used predefined cortical parcellation, HCP-MMP1.0 combined atlas (shown in the inset at the bottom of Fig. 3c and 4c) from HCP-MMP1.0 (Human Connectome Project Multi-Modal Parcellation version 1.0) (Glasser et al., 2016). The HCP-MMP1.0 atlas provides 180 neocortical areas (360 areas in both hemispheres; see Supplementary Figure 1 in Glasser et al. (2016)). The combined atlas provides 22 sections in each hemisphere (44 in both hemispheres) from the original map. The size of the regions of interest is large; however, we used it to reduce computational complexity.

### Speech data processing

#### Auditory speech signal processing

The amplitude envelope of auditory speech signals was computed following the approach introduced in Chandrasekaran et al. (2009). We first constructed eight frequency bands in the range of 100-10,000 Hz to be equidistant on the cochlear map (Smith et al., 2002). The raw auditory sound speech signals were band-pass filtered in these frequency bands using a fourth-order forward and reverse Butterworth filter. Then Hilbert transform was applied to obtain amplitude envelopes for each band of the signal. The envelopes were then averaged across bands resulting in a wideband amplitude envelope. The final signals were resampled to 250 Hz for further analysis to match the sampling rate of preprocessed MEG data.

#### Visual speech signal processing

Lip area information over time was computed from the speaker’s lip in each video using an in-house MATLAB script as reported in our previous publication (Park et al., 2016) as a visual speech signal for each talk. We first extracted the lip contour of the speaker frame-by-frame and computed lip areas which were then resampled to 250 Hz to match the sampling rate of the preprocessed MEG signal and auditory sound envelope signal.

#### Segmentation of auditory speech

In order to apply the topic model (see below) to the current speech materials, we first segmented speech text from each talk into multiple chunks based on their acoustic properties. Using the Syllable Nuclei (de Jong and Wempe, 2009) in Praat software (Boersma and Weenink, 2018), we obtained acoustic chunks with parameters of 0.25 s minimum pause (silence) durations, -25 dB silence threshold, and 1 s minimum duration of each speech chunk (Figure 1a in Park and Gross (2022)). The segmentation resulted in 129 speech chunks on average with a mean duration of 3.37 s from the seven talks used for auditory speech talks.

#### Speech text processing and LDA topic model

Transcriptions of TED talks were downloaded and used for video filming. After the video filming by a professional speaker, the transcriptions were updated as to the differences in the actual speech during the filming. Annotations from the segmentation steps using the Syllable Nuclei were yielded to the text preprocessing step – Tokenization, lemmatization, and removal of stop words for topic model application at a later stage. The preprocessing of text materials was performed using the spaCy (Honnibal and Montani, 2017), an open-source library for Natural Language Processing (NLP) and Scikit-learn (Pedregosa et al., 2011). In spaCy, an English language model, OntoNotes 5 (Weischedel et al., 2013) by Linguistic Data Consortium (LDC), which was trained on a large corpus comprising various genres of text (news, conversational telephone speech, weblogs, usenet newsgroups, broadcast, talk shows), is used. First, raw speech texts were tokenized into component pieces, e.g., prefix, suffix, infix, and other exceptions. Then the tokenized components were lemmatized, e.g., “was” to “be”. Finally, semantically uninformative words (stop words) such as “a/an”, “the”, “be-verbs”, and “and” etc. were filtered out from the text.

Next, the Latent Dirichlet Allocation (LDA) topic model (Blei et al., 2003; Blei, 2012) was applied to the preprocessed text using Scikit-learn (class: sklearn.decomposition.LatentDirichletAllocation). LDA is a generative probabilistic model for collections of texts that implements a three-level hierarchical Bayesian model. The intuition behind the LDA model is that documents are probability distribution over a set of latent topics, and topics themselves are probability distribution over words. The LDA algorithm is typically applied to different documents to classify the documents according to their topics. In the current study, we used the LDA to derive topic keywords that best represent the main idea of each talk across the segmented speech chunks described above.

For the topic model, we used bi-gram (two words) model since a phrase represents the main idea of the talk, such as “machine age”, not “machine” or “age” alone. LDA requires setting k number of topics for fitting the model, which assumes k number of topics are present over multiple documents. In the current study, we assumed four topics are present over segmented speech chunks of each talk. For more details on text preprocessing and topic model application, please see Materials and Methods in Park and Gross (2022). We here have reprinted it below.

For every word (bi-gram in the current study) in every document (speech chunk in the current study) and for each topic:

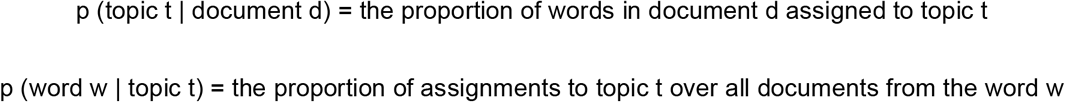

Then the model reassigns w a new topic where topic t with probability:

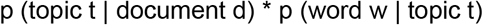

that represents the probability that topic t generated word w.

After applying the model, documents assigned to topic *t* and words with a high probability for topic *t* are obtained as outputs as a document-topic matrix. Also, the most common words (bi-grams in the current study) with the highest probability for topic *t* can be obtained, as shown in Fig. 1 in Park and Gross (2022).

#### Representational interactions between information quantities in the neural processing of auditory and visual speech

Information theory, which was initially developed for studying communications systems (Shannon, 1948), provides a theoretical and practical framework for statistical analysis in neuroscience, including MEG times series data (Gross et al., 2013; Ince et al., 2015; Park et al., 2015; Park et al., 2018b; Park et al., 2018a). Mutual information (MI), one of the quantities in information theory, is a measure of statistical dependence between two variables with a meaningful effect size measured in bits. MI of 1 bit corresponds to a reduction of uncertainty about one variable of a factor 2 after observation of another variable. We previously used MI as a metric for the computation of brain activities entrained by external signals such as speech particularly using Gaussian-Copula Mutual Information (GCMI) (Ince et al., 2017) which implements a robust, semiparametric lower bound estimator of MI by combining the statistical theory of copulas with the closed-form solution for the entropy of Gaussian variables. The advantage of this method is that it performs well for higher dimensional responses as required for measuring three-way statistical interactions and allows estimation over circular variables, like phase information in oscillatory activities.

#### Interaction Information (II) theory

In the previous literature, we showed the representational interactions between dynamic audio and visual speech signals using the information-theoretic approach, Partial Information Decomposition (PID) (Park et al., 2018b). For more detailed theoretical and technical background regarding the PID approach, please see (Williams and Beer, 2010; Ince, 2017; Wibral et al., 2017). In the current study, we used a similar approach, Interaction Information (McGill, 1954) (also known as co-information (Bell, 2003)), that allows us to determine different types of representational interactions between auditory and visual speech information in the brain; redundancy and synergy. Moving beyond the univariate mutual information (MI) analysis, we seek to study the relationship between the neural representations of multi-modal tasks (e.g., auditory and visual speech perception) or distinct spatial regions (e.g., different brain regions) or spectral regions (e.g., alpha and gamma), imaging modalities (e.g., fMRI and M/EEG) (Ince et al., 2017). Interaction Information (II) or PID provides not only redundant (similar or shared) information representation but also synergistic information representation from those relationships (Schneidman et al., 2003; Panzeri et al., 2008; Timme et al., 2014).

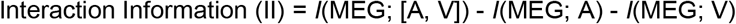

wherein positive II represents synergy and negative II represents redundancy. Please note that this is equivalent to the co-information but with the opposite sign (Bell, 2003). In this study, the redundancy quantifies the shared mutual information (MI) between neural responses to auditory and visual speech, whereas the synergy quantifies the MI in the pair of neural responses to auditory and visual speech when considered jointly is greater than the MI when they are considered individually.

#### Interaction Information (II) analysis

Frequency-specific (1 to 12 Hz in steps of 1 Hz) brain activation time series were computed by applying the Dynamical Imaging of Coherent Sources (DICS) beamformer (Gross et al., 2001) coefficients to the MEG data filtered in the same frequency band (fourth-order Butterworth filter, forward and reverse, center frequency ± 2 Hz). The auditory and visual speech signals were filtered in the same frequency band. MEG signals were shifted by 100 ms as in previous studies to compensate for conduction delays between speech presentation and brain responses (Gross et al., 2013; Park et al., 2016). Redundancy and synergy maps were computed using these auditory and visual speech signals and source-localized brain signals for each voxel and each frequency. From the Hilbert transform, the complex spectra were obtained and normalized by the amplitude. The real and imaginary parts were each rank-normalized. The covariance matrix of the full 6-dimensional signal space was then computed, which completely describes the Gaussian-Copula dependence between the variables. The Interaction Information calculation was performed independently for each voxel and each frequency, resulting in volumetric maps for the three information quantities: joint information, *I*(MEG; A, V), MI between MEG and auditory speech, *I*(MEG; A), and MI between MEG and visual speech, *I*(MEG; V). The analysis was performed for each epoch in each condition and averaged over epochs within the condition (high and low topic probability) and over the delta, theta, and alpha frequency band (1-3 Hz for delta, 4-7 Hz for theta).

#### Conditional Mutual Information (CMI) for each modality speech

To determine modality-specific information representation for each auditory and visual speech, we quantified the conditional mutual information (CMI) from Interaction Information (II) computation described above. Conditional Mutual Information (CMI) is analogous to partial correlation, which quantifies the relationship between two variables while controlling the value of a third variable (Cover and Thomas, 1991; Ince et al., 2012; Ince et al., 2017). Here we calculated the CMI between MEG response and auditory speech conditioning out the effect of visual speech, *I*(MEG; A|V) and the CMI between MEG response and visual speech conditioning out the effect of auditory speech, *I*(MEG; V|A) from the II computation in the following methods.

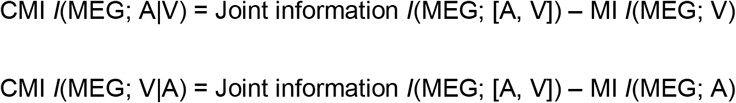

Information-theoretic analyses described in this section (II and CMI) were performed using the Gaussian-Copula Mutual Information (GCMI) toolbox (https://github.com/robince/gcmi) (Ince et al., 2017).

#### Statistical test

For statistical analyses for conditional mutual information (CMI) and Interaction Information (II), the non-parametric cluster-level paired permutation test based on a t-statistic (Maris and Oostenveld, 2007) was performed (function: mne.stats.permutation_cluster_1samp_test) between high versus low topic probability conditions after morphing into common cortical space (fsaverage) in MNE-Python (Gramfort et al., 2013). Cluster-level permutation test handles multiple comparison problems with cluster-level correction and only results at a significance level p < 0.05 are reported. A spatial adjacency matrix was used for clustering in source space, and 1024 permutations were computed.

In order to determine redundant and synergistic brain regions predicted by behavioural performance, ordinary least square linear regression analysis was performed on the redundant and synergistic maps with speech comprehension accuracy as a regressor across participants in MNE-Python (function: mne.stats.linear_regression), and results are reported after multiple comparison correction was performed using False Discovery Rate (FDR; *p* < 0.05) (Genovese et al., 2002).

Two-way ANOVA analysis with factors of attention (All congruent and AV congruent) and topic probability (high and low probability) was performed using the Python module statsmodels (function: statsmodels.regression.linear_model.OLS, statsmodels.stats.anova.anova_lm) and results at a significance level p < 0.05 are reported.

## Discussion

Our previous report demonstrated brain activities involved in the understanding of topic keywords during natural speech perception using a computational topic modelling algorithm in combination with the encoding and decoding model approach (Park and Gross, 2022). Speech chunks were first derived based on acoustic properties using silence threshold in continuous speech, which can be considered as perceptual chunks above phrasal level. Then Latent Dirichlet Allocation (LDA) topic model was applied to the speech chunks, and corresponding brain activities were investigated. This allows us to investigate higher-level semantic processing beyond lexico-semantics which has been unknown in the field. In the current study, we aimed to identify the interaction between auditory-visual information modulated by slow frequency brain activities for topic keyword processing in a multi-speaker environment.

The main findings can be summarized as follows: First, synergistic interaction was represented in delta phase information, whereas redundant interaction between auditory and visual speech was represented in theta phase information in a multi-speaker environment. Second, both types of representational interactions were stronger for speech chunks with low topic probability. This was supported by behavioural performance (by regression analysis using speech comprehension scores in Fig. 3c and Fig. 4c) and anatomical connectivity (by correlation with volumes of white matter fiber tracts obtained using probabilistic tractography analysis in Fig. 5). Third, when compared to the single speaker condition, the interaction information provides deeper insights into the functional role of delta and theta frequency bands in processing semantic gist in a multi-speaker condition. The results show that greater interactions for low semantic gist are represented in synergy by delta, which is sensitive to the speaker condition and redundancy by theta, which is more optimized for tracking semantic contents.

Note that greater positive and negative values represent stronger synergistic and redundant interactions representation. Conceptually, not computationally, the synergistic and redundant interactions are analogous to the supra-additive effect and conjunction effect, respectively (see Discussion in Park et al. (2018b)). The greater Interaction Information for low compared to high topic probability condition for both types of Interaction Information processing and their behavioural relevance supports the interactions between two modalities (auditory and visual) are more required for processing speech with less semantic gist. To be able to obtain a high score on the behavioural speech comprehension tests, exhaustive attention throughout the speech perception might have been required. Specifically, during speech perception in a multi-speaker situation, it might entail more effortful listening, and a compensational mechanism is needed for speech chunks which are semantically less salient, i.e., low topic probability condition in the current study. This can be interpreted in the light of efficient information processing in the brain. It has been known that optimization of information processing in the human brain relies on both strong and weak connections (Schwarz and McGonigle, 2011; Gallos et al., 2012), and engagement of multiple distributed networks is evident during effortful tasks compared to effortless tasks (Finc et al., 2017). Thus, less semantically salient speech chunks which require more attention and intelligibility for comprehension processing demand more distributed brain engagement. The significantly stronger correlation between whiter matter fiber tract volumes and Interaction Information measures for low semantic gist from both delta synergy and theta redundancy also support this notion that subjects with larger volumes of the whiter matter tracts exert more efficient interactions between audiovisual modalities. Particularly, the theta redundancy in the right temporo-parieto-occipital junction shows significant relationships with all the major tracts. This region is known to be a complex territory connecting temporal, parietal and occipital areas involving high-level cognitive functions such as multi-modal integration, attentional control, language, reading, writing, memory, visual, and social cognition etc. (Duffau, 2008; Geng and Mangun, 2011; De Benedictis et al., 2014). A recent study has shown that this region is covered by thick white matter connections, which provide efficient information processing as a crucial node for cortical and subcortical connectivity (De Benedictis et al., 2014). Our results provide evidence that this region supports more efficient redundant interactions for less semantically salient speech processing by theta phase information.

Since our findings emerged in the multi-speaker environment, we sought to compare the dynamics by comparing the single-speaker condition (natural audiovisual speech condition) using two-way ANOVA with the factors of speaker condition and topic probability (Fig. 6). We found three interesting patterns: 1) the Interaction Information values of the delta band phase in the left auditory, temporal, and inferior frontal areas were observed for the main effect of speaker condition, whereas 2) the same regions in the right hemisphere were observed for the interaction effect of speaker condition and topic probability, and 3) the Interaction Information values of theta phase in right auditory, temporal and bilateral temporal, frontal areas were observed for the main effect of topic probability. Furthermore, similar patterns were observed in other areas as well as shown in Supplementary Figure 1.

Synergistic information processing in the delta band in the left regions has shown the main effect of speaker condition, suggesting that the delta phase information in the left auditory, temporal and inferior frontal regions are sensitive to the speaker environment. The results show that synergistic information by delta phase was greater for the single speaker condition than for the multi-speaker condition. However, in the single speaker condition, there is no difference between high and low topic probability conditions, unlike in the multi-speaker condition. Interestingly, the same regions in the right hemisphere have shown an interaction effect of speaker condition and topic probability. In the single speaker condition, synergistic information was greater for the high topic probability condition; however, this pattern is reversed in the multi-speaker condition. This suggests that the sensitivity to high versus low semantic gist exerting greater synergistic information out of the two modalities depends on the speaker environment in the right hemisphere regions. The hemispheric difference and lateralization for specific speech features in speech perception have been one of the key topics in the field (Poeppel, 2003; Gross et al., 2013; Flinker et al., 2019; Giroud et al., 2020). In the context of semantic processing, the right hemisphere is known to be engaged in semantic control (Bowden and Beeman, 1998; Thompson et al., 2016) and drawing coherent inferences using semantic information (Beeman, 1993). Furthermore, given the complexity in the processing of higher-level semantic gist during naturalistic speech perception, it is assumed that not only linguistic features but higher-level feature processing, e.g., connotations, pragmatics etc., are involved. In such processing, the right hemisphere activation has been suggested to involve in coherence and cohesion in an effort to integrate meanings and is concerned with deriving the general idea from the key message through complex and abstract linguistic interpretation (Beeman, 1993; Campbell, 2006). This processing becomes more sensitive for semantically less salient speech in a multi-speaker environment. Meanwhile, redundant information by theta band in these brain regions in both hemispheres has shown the main effect of topic probability. Stronger redundancy (more negative values) for low semantic gist processing in both single- and multi-speaker conditions was observed though the findings in left early and associative auditory areas should be considered as a trend given statistically non-significant results (but see other areas in Supplementary Figure 1). This suggests that the shared mechanism (redundancy) out of the two modalities carried by the theta band is sensitive only to semantic saliency regardless of the speaker environment.

Higher-order semantic processing in natural speech perception is complex processing indeed integrating sensory, short- and long-term memory processing, interpreting social context, deriving meaning and making inferences. It becomes even more complicated in multi-modal and multi-speaker environments. The current study demonstrated neural oscillatory mechanism in representational interactions of multi-modal speech features for topic keyword processing and provided insights into the differential role in delta and theta bands as well as hemispheric asymmetry modulated by delta phase information. Particularly, speaker environment dependant processing for topic saliency could be further investigated in wider naturalistic speech perception settings by extending from dichotic listening to more realistic cocktail party-like situations and discourse-level communication.

## Data availability

The data that support the findings of this study are available from the corresponding author upon request.

## Code availability

The computer code and algorithms used to generate our results are available from the corresponding author upon request.

## Acknowledgements

Funding: Wellcome Trust Senior Investigator Award (grant number: 098433) and DFG project funding (GR 2024/5-1, GR 2024/11-1) to JG. Wellcome Trust Seed Awards in Science (grant number: 214120/Z/18/Z) to RAAI. The funder had no role in study design, data collection and analysis, decision to publish, or preparation of the manuscript.

## Author contributions

Conceptualization: H.P., J.G.; Data collection: H.P.; Data analysis: H.P.; Methodology: H.P., R.A.A.I., J.G.; Project administration: H.P., J.G.; Writing: H.P., R.A.A.I., J.G.

## Conflict of interest statement

The authors declare that the research was conducted in the absence of any commercial or financial relationships that could be construed as a potential conflict of interest.

## Notes

### Competing Interest Statement

The authors have declared no competing interest.

