## Supplementary Information for "Differential roles of delta and theta oscillations in understanding semantic gist during natural audiovisual speech perception: Functional and anatomical evidence"

**Supplementary Figure 1 (related to Fig. 6).** Two-way analysis of variance (ANOVA) test with factors of speaker condition (single versus multi-speaker; All congruent versus AV congruent conditions) and topic probability (high versus low topic probability) was performed using the Interaction Information for each delta and theta phase information. Here we used the predefined cortical parcellation, HCP-MMP1.0 combined atlas, and the results below contain the regions shown in Figure 6. HCP-MMP1.0 combined atlas provides 22 sections in each hemisphere (44 regions in both hemispheres). We here show all the regions except the following six regions: Anterior Cingulate and Medial Prefrontal Cortex, MT+ Complex and Neighboring Visual Areas, Orbital and Polar Frontal Cortex, Paracentral Lobular and Mid Cingulate Cortex, Posterior Cingulate Cortex and Superior Parietal Cortex, which are more medial brain regions or did not show significant results. *Abbreviations*. *LH*: left hemisphere; *RH*, right hemisphere; *S*, the main effect of speaker condition; *T*, the main effect of topic probability; *I*: the interaction effect between speaker condition and topic probability.

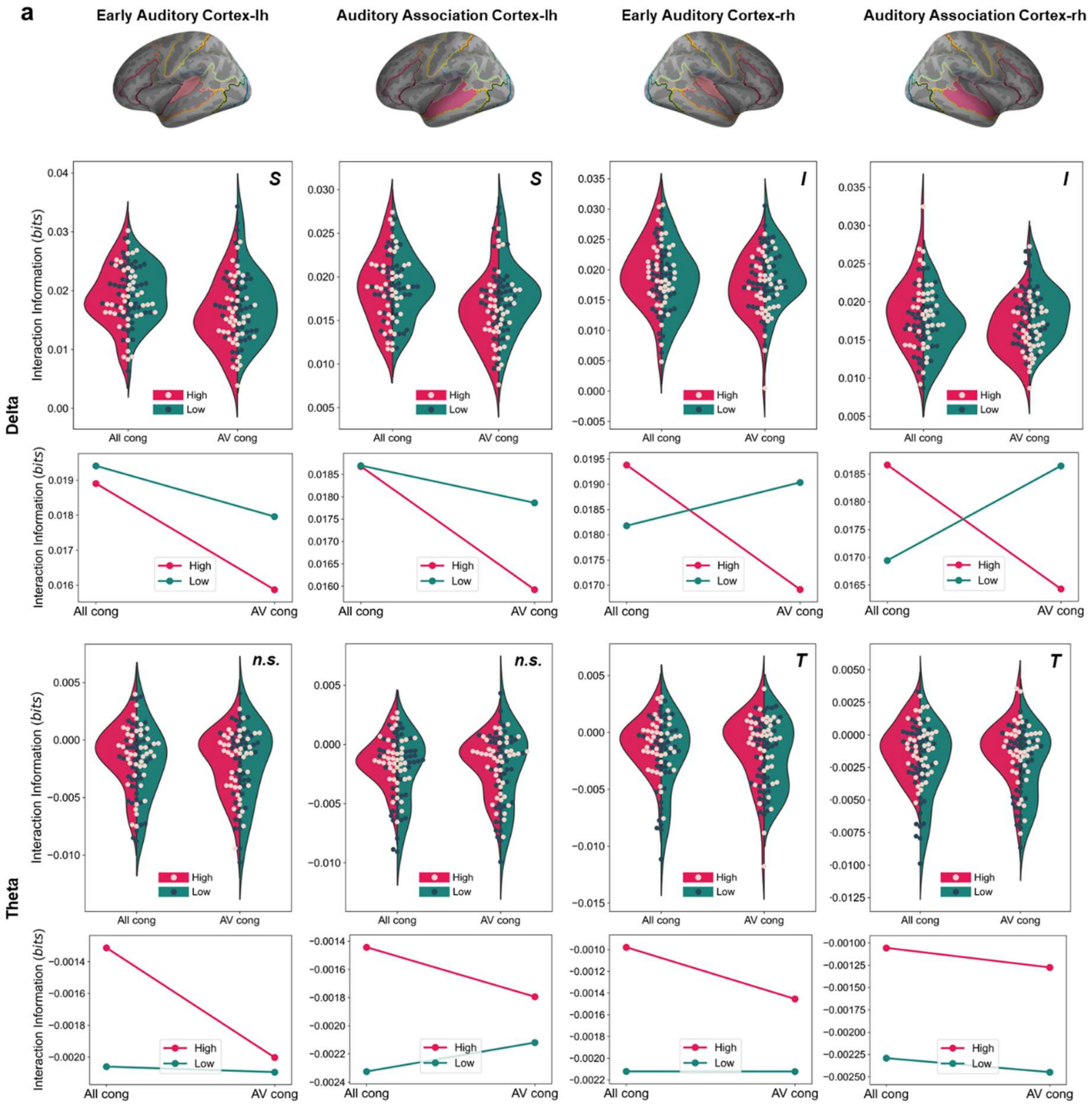


**a, Delta band in the LH** (left two columns in top row): Early auditory cortex (*S*: $F_{1,172} = 7.47$, $p = 0.006$), Auditory association cortex (*S*: $F_{1,172} = 9.26$, $p = 0.002$). **Delta band in the RH** (right two columns in top row): Early auditory cortex (*I*: $F_{1,172} = 4.46$, $p = 0.03$), Auditory association cortex (*I*: $F_{1,172} = 11.12$, $p = 0.001$). **Theta band in the LH** (left two columns in bottom row): Early auditory cortex (all *n.s.*), Auditory association cortex (all *n.s.*). **Theta band in the RH** (right two columns in bottom row): Early auditory cortex (*T*: $F_{1,172} = 4.16$, $p = 0.04$), Auditory association cortex (*T*: $F_{1,172} = 10.20$, $p = 0.001$).

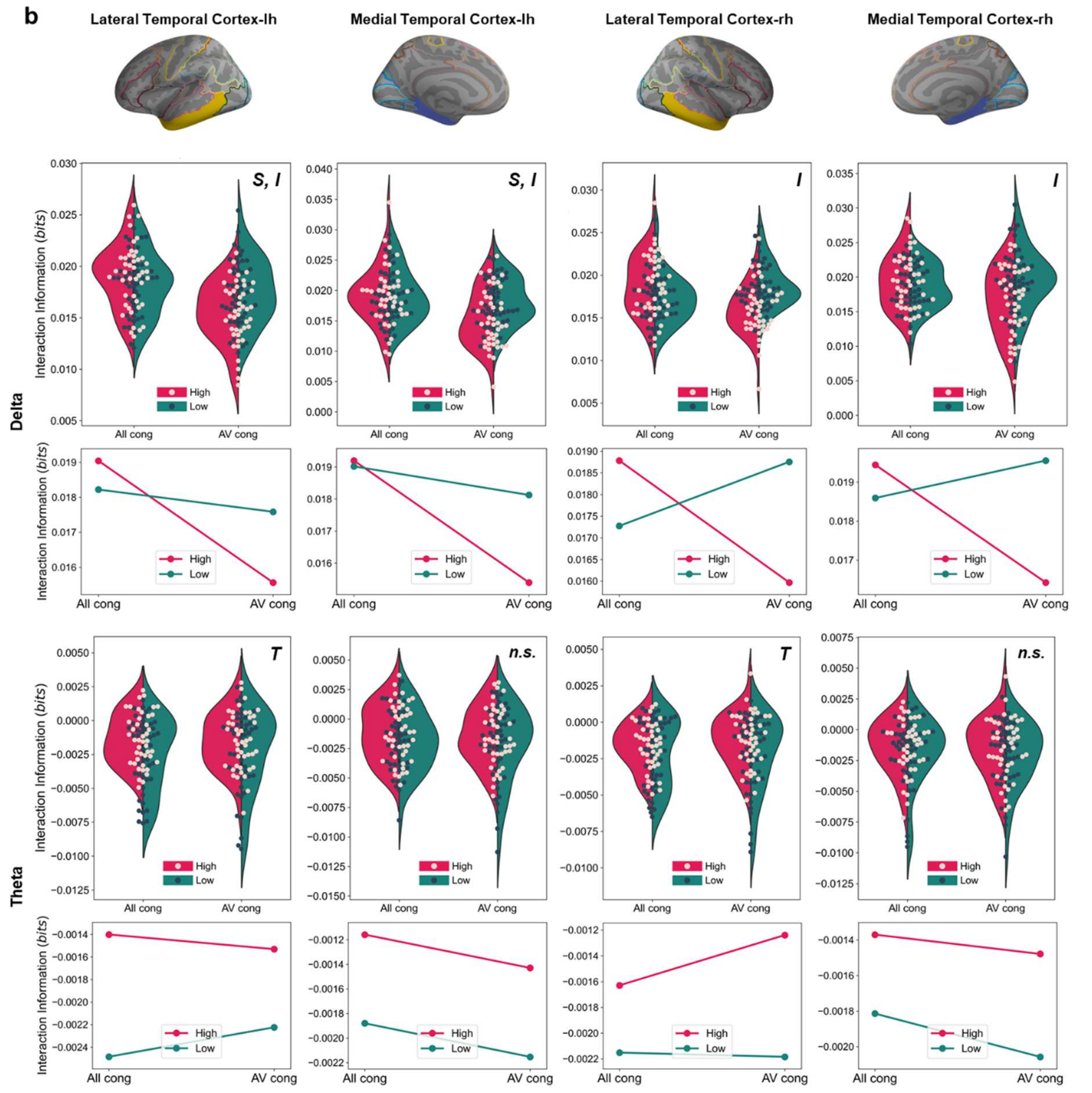


**b, Delta band in the LH** (left two columns in top row): Lateral temporal cortex (*S*: $F_{1,172} = 19.38$, $p < 0.001$; *I*: $F_{1,172} = 9.26$, $p = 0.002$), Medial temporal cortex (*S*: $F_{1,172} = 12.25$, $p < 0.001$; *I*: $F_{1,172} = 4.71$, $p = 0.03$). **Delta band in the RH** (right two columns in top row): Lateral temporal cortex (*I*: $F_{1,172} = 21.69$ $p < 0.001$), Medial temporal cortex (*I*: $F_{1,172} = 9.71$, $p = 0.002$). **Theta band in the LH** (left two columns in bottom row): Lateral temporal cortex (*T*: $F_{1,172} = 5.67$, $p = 0.01$), Medial temporal cortex (all *n.s.*). **Theta band in the RH** (right two columns in bottom row): Lateral temporal cortex (*T*: $F_{1,172} = 5.15$, $p = 0.02$), Medial temporal cortex (all *n.s.*).

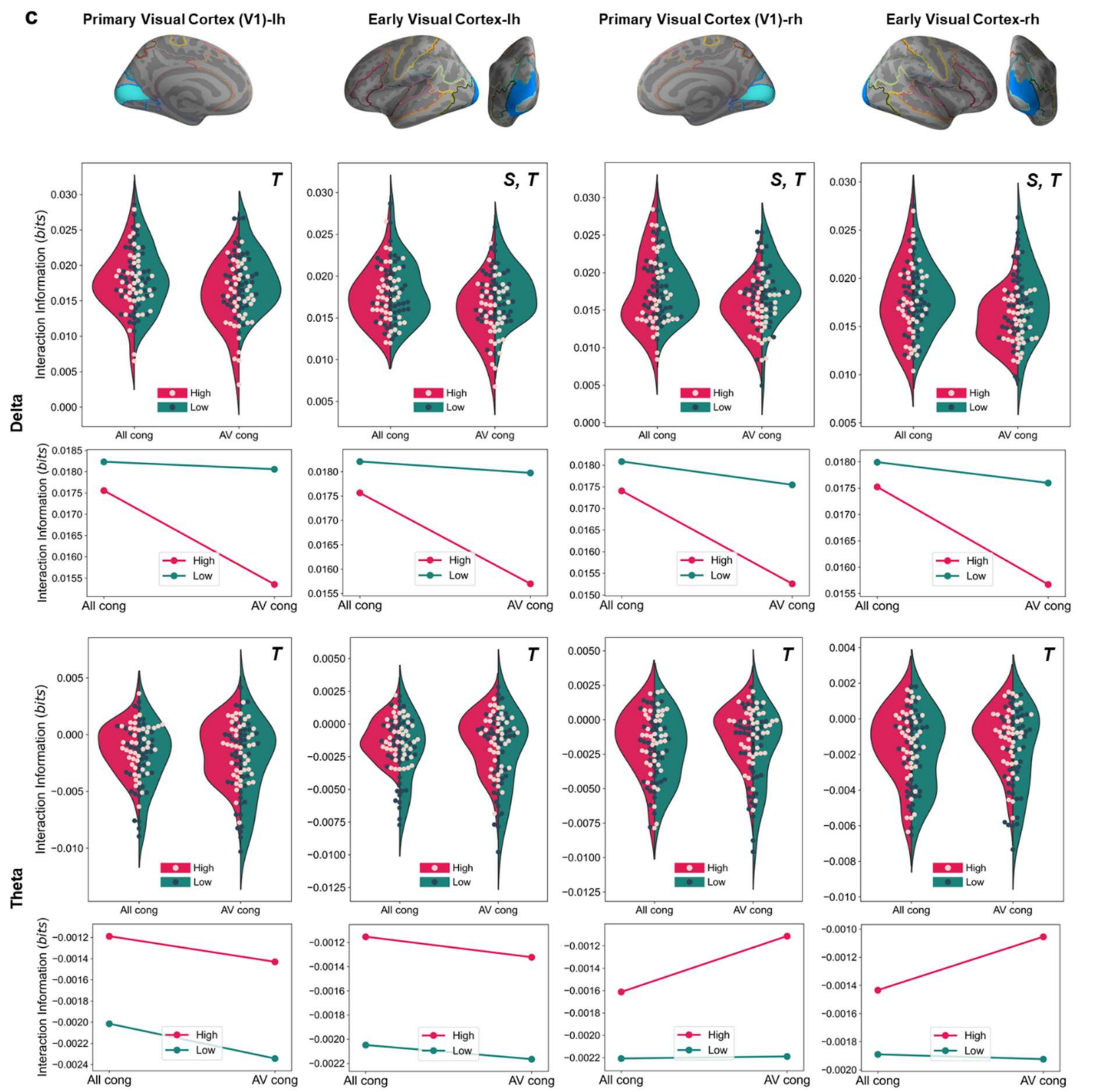


**c, Delta band in the LH** (left two columns in top row): Primary visual cortex (*T*: $F_{1,172} = 6.75$, p = 0.01), Early visual cortex (*S*: $F_{1,172} = 4.11$, p = 0.04; *T*: $F_{1,172} = 7.95$, p = 0.005). **Delta band in the RH** (right two columns in top row): Primary visual cortex (*S*: $F_{1,172} = 4.68$, p = 0.03; *T*: $F_{1,172} = 5.71$, p = 0.01), Early visual cortex (*S*: $F_{1,172} = 4.61$, p = 0.03; *T*: $F_{1,172} = 5.26$, p = 0.02). **Theta band in the LH** (left two columns in bottom row): Primary visual cortex (*T*: $F_{1,172} = 4.25$, p = 0.04), Early visual cortex (*T*: $F_{1,172} = 6.01$, p = 0.01). **Theta band in the RH** (right two columns in bottom row): Primary visual cortex (*T*: $F_{1,172} = 5.07$, p = 0.02), Early visual cortex (*T*: $F_{1,172} = 4.46$, p = 0.03).

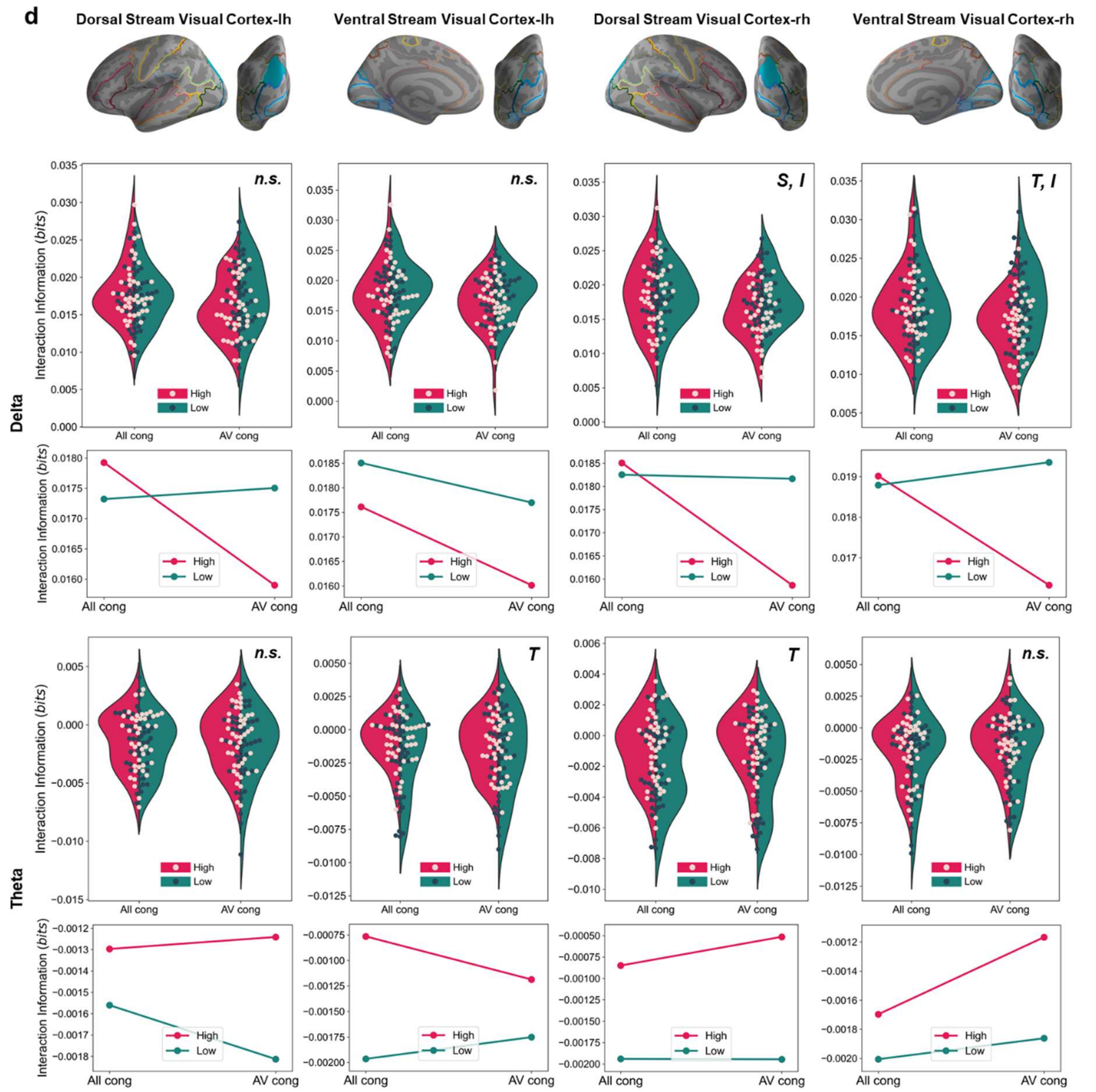


**d, Delta band in the LH** (left two columns in top row): Dorsal stream visual cortex (all *n.s.*), Ventral stream visual cortex (all *n.s.*). **Delta band in the RH** (right two columns in top row): Dorsal stream visual cortex (*S*: $F_{1,172} = 4.33$, $p = 0.03$; *I*: $F_{1,172} = 3.80$, $p = 0.05$), Ventral stream visual cortex (*T*: $F_{1,172} = 4.15$, $p = 0.04$; *I*: $F_{1,172} = 5.59$, $p = 0.01$). **Theta band in the LH** (left two columns in bottom row): Dorsal stream visual cortex (all *n.s.*), Ventral stream visual cortex (*T*: $F_{1,172} = 5.02$, $p = 0.02$). **Theta band in the RH** (right two columns in bottom row): Dorsal stream visual cortex (*T*: $F_{1,172} = 11.71$, $p < 0.001$), Ventral stream visual cortex (all *n.s.*).

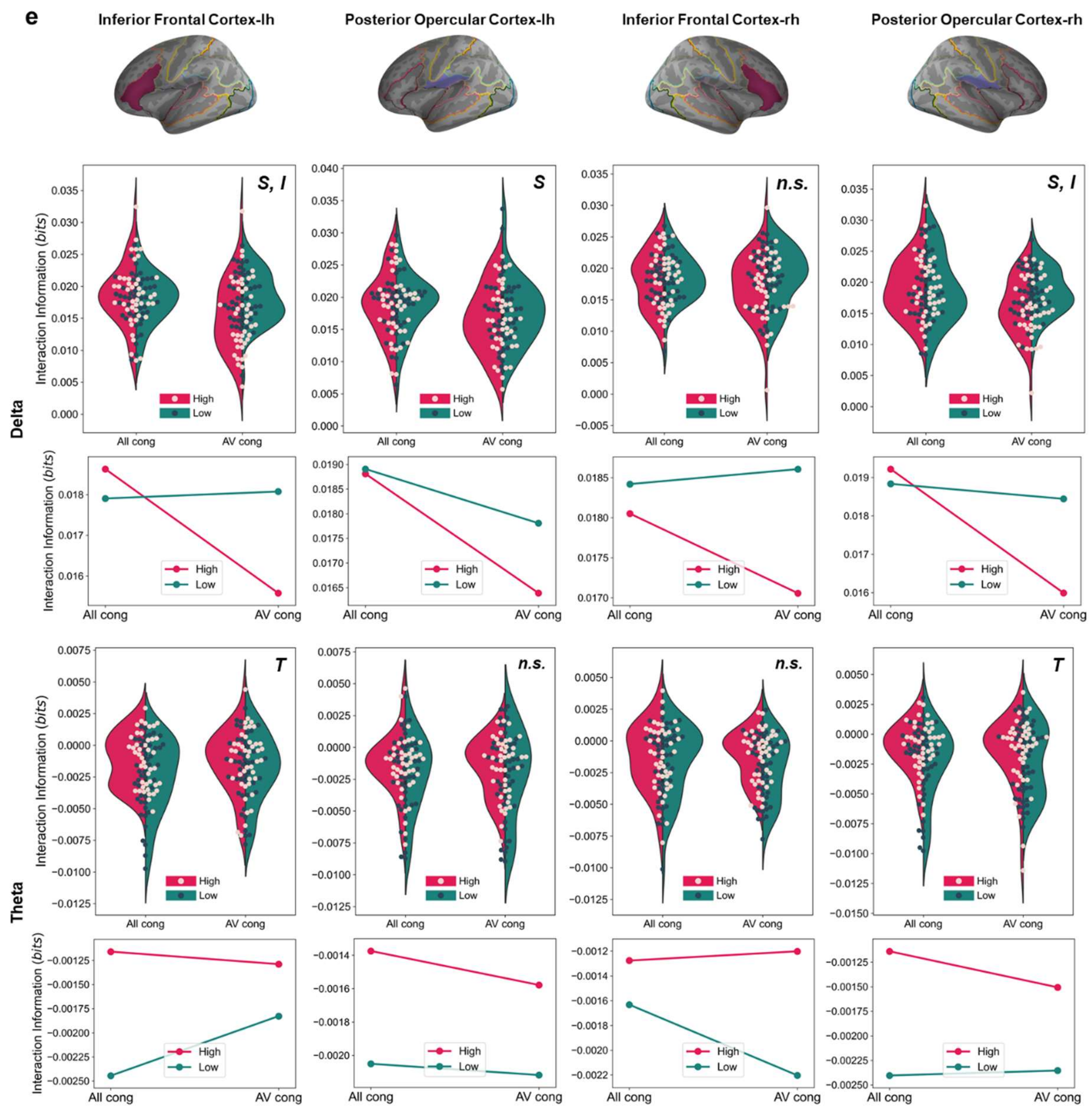


**e, Delta band in the LH** (left two columns in top row): Inferior frontal cortex (*S*: $F_{1,172} = 4.21$, $p = 0.04$; *I*: $F_{1,172} = 5.27$, $p = 0.02$), Posterior opercular cortex (*S*: $F_{1,172} = 5.56$, $p = 0.01$). **Delta band in the RH** (right two columns in top row): Inferior frontal cortex (all *n.s.*), Posterior opercular cortex (*S*: $F_{1,172} = 6.44$, $p = 0.01$; *I*: $F_{1,172} = 3.95$, $p = 0.04$). **Theta band in the LH** (left two columns in bottom row): Inferior frontal cortex (*T*: $F_{1,172} = 5.64$, $p = 0.01$), Posterior opercular cortex (all *n.s.*). **Theta band in the RH** (right two columns in bottom row): Inferior frontal cortex (all *n.s.*), Posterior opercular cortex (*T*: $F_{1,172} = 6.22$, $p = 0.01$).

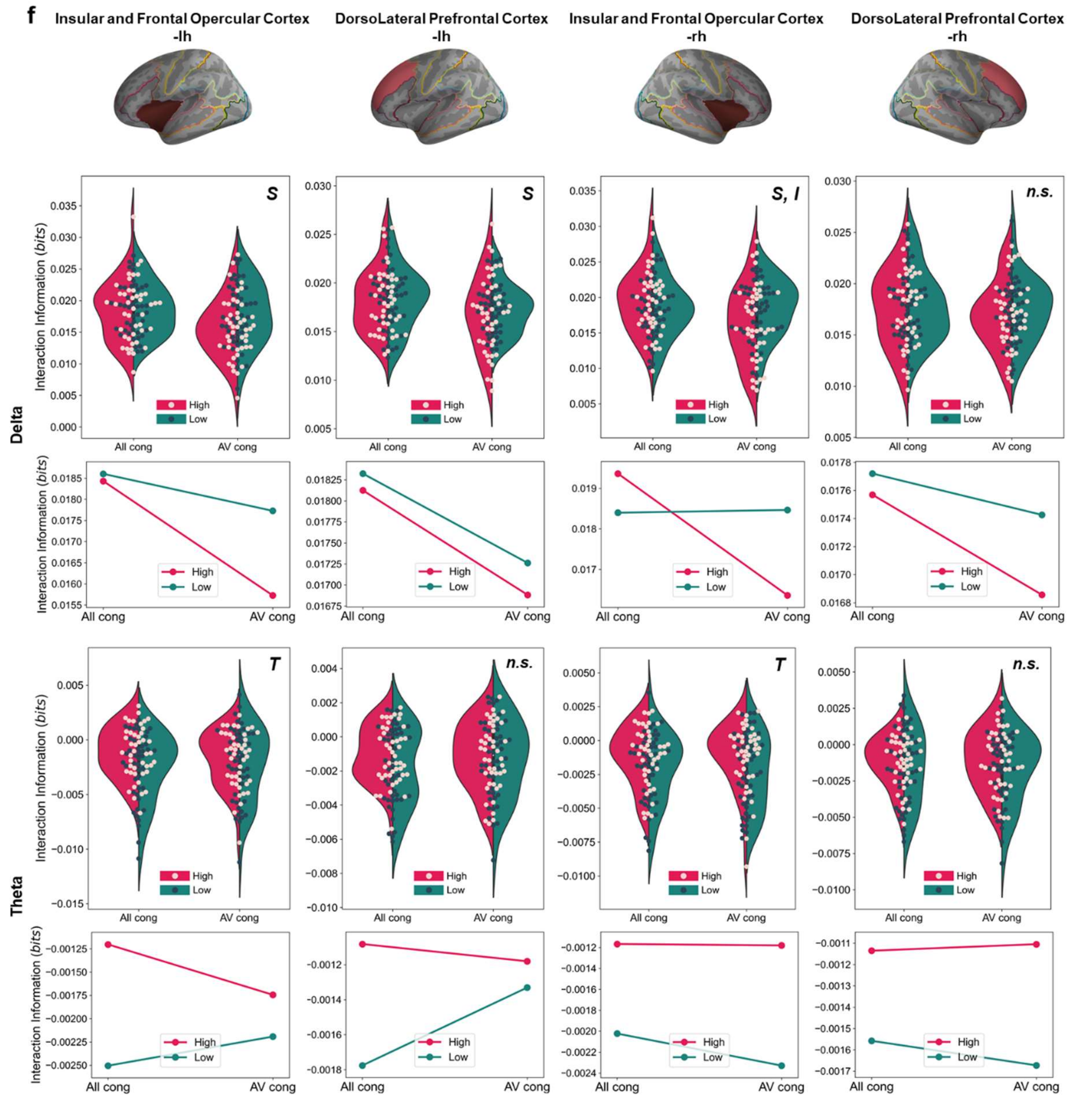


**f, Delta band in the LH** (left two columns in top row): Insular and frontal opercular cortex (*S*: $F_{1,172}$ = 6.44, p = 0.01), Dorsolateral prefrontal cortex (*S*: $F_{1,172}$ = 5.59, p = 0.01). **Delta band in the RH** (right two columns in top row): Insular and frontal opercular cortex (*S*: $F_{1,172}$ = 4.64, p = 0.03; *I*: $F_{1,172}$ = 5.08, p = 0.02), Dorsolateral prefrontal cortex (all *n.s.*). **Theta band in the LH** (left two columns in bottom row): Insular and frontal opercular cortex (*T*: $F_{1,172}$ = 4.00, p = 0.04), Dorsolateral prefrontal cortex (all *n.s.*). **Theta band in the RH** (right two columns in bottom row): Insular and frontal opercular cortex (*T*: $F_{1,172}$ = 7.11, p = 0.008), Dorsolateral prefrontal cortex (all *n.s.*).

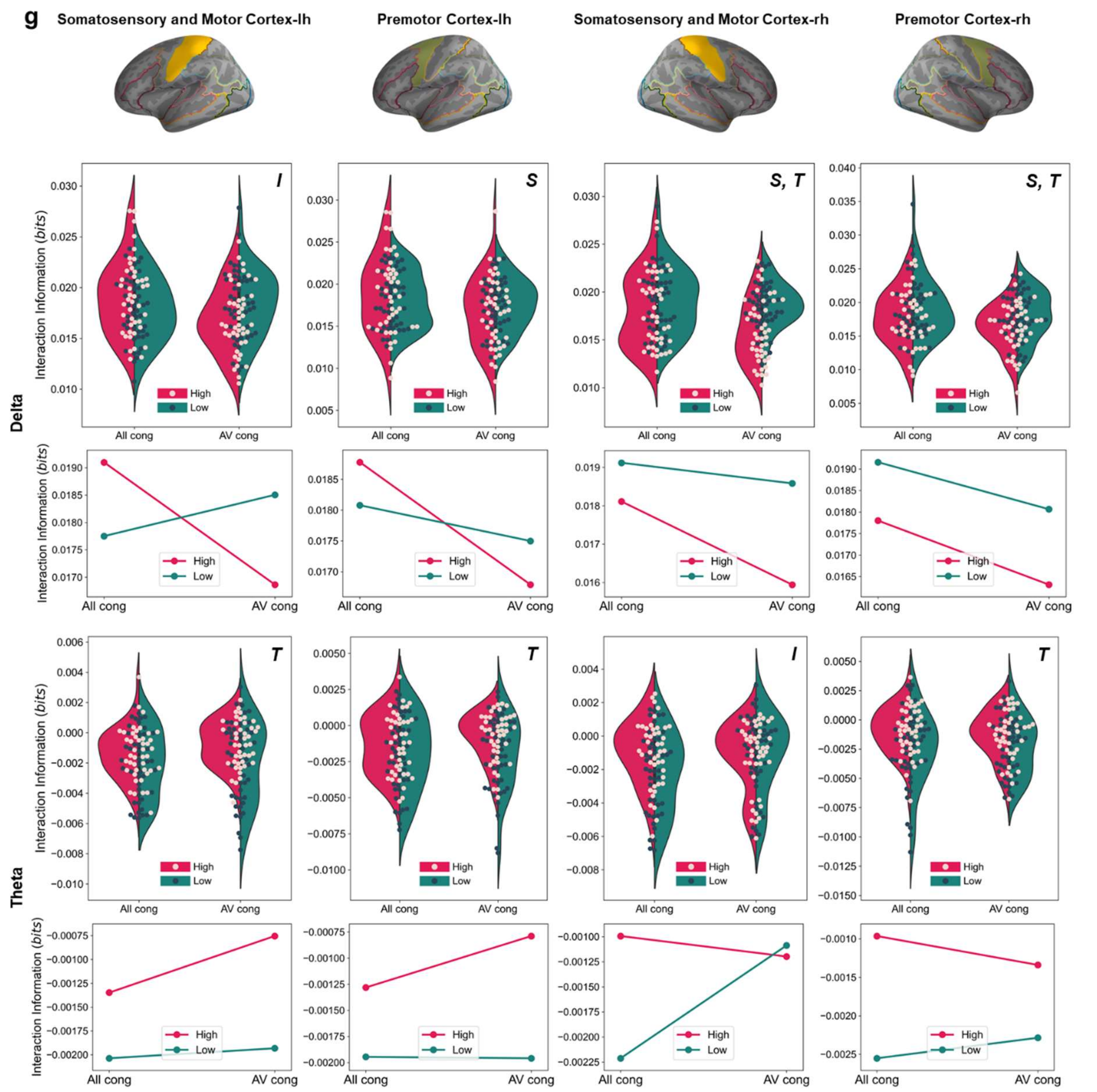


**g, Delta band in the LH** (left two columns in top row): Somatosensory and motor cortex (*I*: $F_{1,172} = 8.52$, $p = 0.003$), Premotor cortex (*S*: $F_{1,172} = 4.94$, $p = 0.02$). **Delta band in the RH** (right two columns in top row): Somatosensory and motor cortex (*S*: $F_{1,172} = 7.39$, $p = 0.007$; *T*: $F_{1,172} = 13.46$, $p < 0.001$), Premotor cortex (*S*: $F_{1,172} = 4.65$, $p = 0.03$; *T*: $F_{1,172} = 6.75$, $p = 0.01$). **Theta band in the LH** (left two columns in bottom row): Somatosensory and motor cortex (*T*: $F_{1,172} = 8.63$, $p = 0.003$), Premotor cortex (*T*: $F_{1,172} = 7.12$, $p = 0.008$). **Theta band in the RH** (right two columns in bottom row): Somatosensory and motor cortex (*I*: $F_{1,172} = 4.02$, $p = 0.04$), Premotor cortex (*T*: $F_{1,172} = 10.94$, $p = 0.001$).

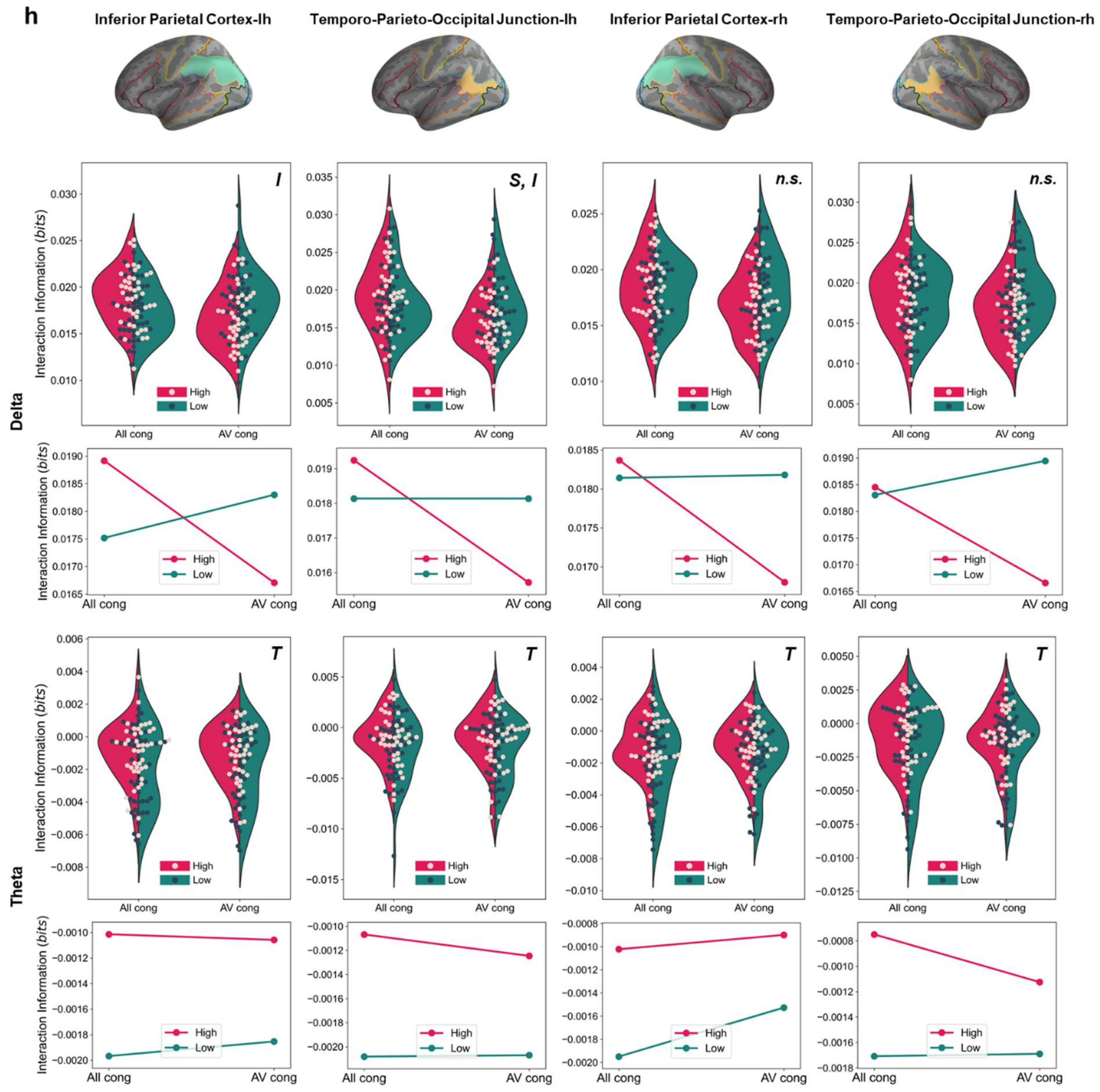


**h, Delta band in the LH** (left two columns in top row): Inferior parietal cortex (*I*: $F_{1,172}$ = 9.90, p = 0.001), Temporo-parieto-occipital junction (*S*: $F_{1,172}$ = 7.47, p = 0.006; *I*: $F_{1,172}$ = 7.46, p = 0.006). **Delta band in the RH** (right two columns in top row): Inferior parietal cortex (all *n.s.*), Temporo-parieto-occipital junction (all *n.s.*). **Theta band in the LH** (left two columns in bottom row): Inferior parietal cortex (*T*: $F_{1,172}$ = 7.30, p = 0.007), Temporo-parieto-occipital junction (*T*: $F_{1,172}$ = 4.34, p = 0.03). **Theta band in the RH** (right two columns in bottom row): Inferior parietal cortex (*T*: $F_{1,172}$ = 6.61, p = 0.01), Temporo-parieto-occipital junction (*T*: $F_{1,172}$ = 3.86, p = 0.05).
